# CPEB2 m^6^A Methylation Regulates BTB Permeability via Affecting Splicing Factor SRSF5 Stability

**DOI:** 10.1101/2020.11.17.387928

**Authors:** Mengyang Zhang, Chunqing Yang, Xiaobai Liu, Xuelei Ruan, Di Wang, Yunhui Liu, Heng Cai, Jian Zheng, Lianqi Shao, Ping Wang, Zhen Li, Bo Yu, Yixue Xue

## Abstract

The existence of the blood tumor barrier (BTB) severely hinders the delivery of anti-tumor drugs to gliomas, affecting the targeted therapeutic effects of drugs. Therefore, BTB selective opening has become a hot spot for glioma treatment. This study found that the up-regulated METTL3 and IGF2BP3 in GECs increase the stability of CPEB2 mRNA through m6A methylation of CPEB2 mRNA; CPEB2 binds and increases the stability of splicing factor SRSF5 mRNA; SRSF5 promotes the ETS1 exon inclusion; P51-ETS1 promotes the transcriptional expression of tight junction related proteins ZO-1, occludin and claudin-5, regulating BTB permeability. CPEB2, SRSF5 and P51-ETS1 alone or in combination can effectively enhance the role of Dox in promoting glioma cell apoptosis through BTB. The results of this study provide a new theoretical and experimental basis for the molecular regulation of BTB from the perspective of epigenetics, as well as new ideas for the comprehensive treatment of glioma.

## Background

Malignant glioma is the most common primary tumor in the central nervous system. Currently the main treatment is surgery-assisted radiotherapy and chemotherapy. Chemotherapy is one of the most commonly used treatments for malignant glioma. Due to the presence of blood tumor barrier (BTB), it is difficult for large-molecule chemotherapy drugs to reach tumor tissues to exert therapeutic effects. Therefore, the selective opening of BTB is an effective strategy to improve the chemical efficacy of glioma. The tight junction composed of tight junction related proteins is one of the main ways to regulate the permeability of BTB. A large number of studies have proved that the reduced expression of TJP, such as the transmembrane proteins occludin and claudin-5, and the cytoplasmic plaque protein ZO-1, can increase BTB permeability^[1]^.

M^6^A is the most abundant methylated modification in eukaryotic mRNA. The m^6^A modification site on mRNA mainly occurs on the adenine of the RRACH (R: purine; A: m^6^A; H: non-guanine) sequence, which is highly conserved^[2]^. The function is determined by methyltransferase (writer), Demethylase (eraser) and binding protein (reader) are jointly regulated^[3]^. In recent years, more and more studies have shown that m^6^A modification can functionally regulate the transcriptome functions of eukaryotes, such as mRNA splicing, nucleation, localization, translation and stability^[3]^. Studies have shown that the modification of m^6^A determines the fate of hematopoietic stem cells during the development of vertebrate embryos^[5]^. METTL3 (methyltransferase like 3) is one of the earliest identified m^6^A methyltransferases. It is highly expressed in a variety of tumor tissues. By regulating the methylation modification of target genes, it promotes mRNA translation and regulates tumor cell proliferation^[6]^. IGF2BP3 (insulin like growth factor 2 mRNA binding protein3) as a member of the m^6^A reading family IGF2BPs (insulin like growth factor 2 mRNA binding proteins), enhances the stability and translation of mRNA by recognizing the GG (m^6^A) C sequence shared by mRNAs^[7]^. Studies have found that IGF2BP3 is highly expressed in lung cancer and breast cancer and regulated the occurrence and development of tumors^[8]^. However, there was no research on IGF2BP3 regulating vascular endothelial function has been retrieved.

RBPs play an important role in regulating gene expression during transcription and post-transcription. RBPs mediate processes such as RNA stability maintenance, intracellular localization and translation regulation by recognizing specific elements of mRNA^[9]^. Recent studies have shown that RBPs are involved in regulating the occurrence and development of many tumors^[10]^. CPEB2 (Cytoplasmic poly (A) denylation element binding protein 2) is a member of the cytoplasmic polyadenylation element binding protein family, which regulates the translation of the target gene mRNA by binding to the cytoplasmic polyadenylation element in the 3’untranslated region^[11]^. The CPEB family inhibits or activates the translation of target gene mRNA by shortening or lengthening the poly-A tail^[12]^. Studies have reported that CPEB2 plays an important role in the occurrence and development of triple-negative breast cancer^[13]^.

As one of the most common gene regulation mechanisms, alternative splicing plays an important role in the complex regulation of protein functions. Splicing dysregulation is closely related to the occurrence and development of various human tumors^[14]^. Alternative splicing of precursor messenger RNA (pre-mRNA) is common in mammalian cells, and nearly 95% of human genes are formed by alternative splicing^[15, 16]^, which helps regulate gene expression and expand proteins diversity of groups. SRSF5 (serine/arginine-rich splicing factor 5) is a member of the serine/arginine-rich protein family, contains two N-terminal RNA recognition motifs (RRM), and an arginine/serine-rich domain^[17]^, play an important regulatory role in RNA splicing and translation. Studies have shown that SRSF5 is abnormally expressed in various tumors such as breast cancer, kidney cancer and lung cancer^[18]^.

ETS1 (ETS proto-oncogene 1, transcription factor) is involved in regulating the proliferation, development, apoptosis, metastasis, invasion and angiogenesis of tumor cells^[19]^. ETS1 can regulate vascular endothelial cell maturation and endothelial barrier function^[20]^. ETS1 contains ETS domain (transcription activation domain) and helical DNA binding domain, and is highly expressed in various tumor tissues^[21]^. Inhibiting ETS1 can block tumor proliferation, migration and invasion in vivo^[22]^. Until now, there have been no research reports on the expression of METTL3, IGF2BP3, CPEB2, SRSF5 and ETS1 in glioma endothelial cells (GECs), and the regulation of BTB permeability.

In this study, we first investigated the endogenous expression of METTL3, IGF2BP3, CPEB2, SRSF5, and ETS1 in GECs, and further analyzed the mode of action between these molecules, as well as the role and molecular mechanism of regulating BTB permeability. This research aims to reveal new mechanisms for regulating BTB permeability, as well as provide new ideas and approaches for selectively opening the blood-brain barrier, increasing drug transport to tumor tissues, and improving the efficacy of chemotherapy.

## Materials and methods

### Cell Lines and Cell Culture

The immortalized human brain endothelial cell line hCMEC/D3 was provided by Dr. Couraud from the Cochin Institute in Paris, France. Endothelial cells were cultured on culture inserts (0.4-mm pore size; Corning, Lowell, MA, USA) coated with 150 μg/mL Cultrex rat collagen I (R&D Systems, Minneapolis, Minnesota, USA). Cells were cultured in endothelial base medium (EBM-2) with 5% fetal bovine serum (FBS) “Gold” (PAA Laboratories, Pasching, Austria), 1% penicillin-streptomycin (Life Technologies, Paisley, UK), 1.4 mol/ L hydrocortisone (Sigma-Aldrich, St Louis, Missouri, USA), 1% chemical determination (Life Technologies, Paisley, UK), 5 g/ mL ascorbic acid (Sigma-Aldrich), 10 mmol/L HEPES (PAA Laboratory) and 1 ng/mL human basic fibroblast growth factor (bFGF) (Sigma-Aldrich) (Lonza, Walkersville, MD, USA). The generation number of ECs is maintained between 30-40 generations. The human glioblastoma U251 cell line and HEK293T cell line were purchased from the Cell Resource Center of Shanghai Institute of Biological Sciences and stored in DMEM which contains 10% FBS, 100 U/mL penicillin and 100 μg/mL streptomycin (Life Technologies, Paisley), United Kingdom). Normal human astrocytes (NHA) were purchased from Sciencell Research Laboratories (Carlsbad, California, USA) and cultured in astrocyte culture medium RPMI-1640 (GIBCO, Carlsbad, California, USA). All cells were cultured in a humidified incubator at 37°C and 5% CO2.

### Establishment of in Vitro BTB and BBB Model

The *in vitro* BTB model was established by co-cultivation of ECs and U251 cells, as we described previously^[54]^. U251 cells were seeded in six-well plates at a density of 2×104 per well and cultured for 2 days. Endothelial cells were seeded at a density of 2×105 per well on the insert coated with Cultrex rat collagen I (R&D Systems, Minneapolis, Minnesota, USA), and the insert was placed in the above-mentioned six-well plates. After co-culturing for 4 days, GECs were obtained. Both endothelial cells and U251 cells were cultured with the prepared EBM-2 medium, and the medium was changed every 2 days. The *in vitro* BBB model used the same method to co-culture endothelial cells with NHA to obtain AECs.

### qRT-PCR Assay

Total RNAs were extracted by Trizol reagent (Life Technologies, Carlsbad, CA, USA). SYBR PrimeScript primary RT-PCR kit (Takara Bio, Japan) was used to evaluate the expression levels of METTL3, IGF2BP3, CPEB2, SRSF5 and ETS1, the reaction was carried out by 7500 Fast RT-PCR system (Applied Biosystems, Foster City, CA, USA). Endogenous control was glyceraldehyde 3-phosphate dehydrogenase (GAPDH). Relative expression values were calculated using the relative quantifification (2^-△△Ct^) method. The primers are provided in Table S1.

### Cell Transfection

Cell transfections were performed as previously described^[55]^. Endothelial cells were seeded in 24-well plates and transfected using Opti-MEM I and Lipofectamine LTX reagent (Life Technologies, Carlsbad, CA, USA) under approximately 80% fusion conditions according to the instructions. Stable cell lines were selected by geneticin (G418) or purinomycin. After 4 weeks of application, G418-resistant (or purinomycin-resistant) clones were obtained. Plasmids and corresponding empty vectors are constructed by the GenePharma (Shanghai, China). The sequences and vectors of plasmids are shown in Table S2.

### Transendothelial electric resistance (TEER) assays

The TEER value was measured using Millicell-ERS instrument (Millipore, Billerica, Mass) after establishing the *in vitro* BTB model. Replace the same amount of new mediums in the upper and lower chambers of Transwell, and measure after 30 min at room temperature. By subtracting the background resistance from the measured blocking resistance and then multiplying it by the effective surface area of the filter was the final resistance (Ωcm^2^).

### Horseradish peroxidase flux (HRP) assays

After establishing the *in vitro* BTB model, added 0.5 µmol/L horseradish peroxidase to upper chamber of Transwell, and incubated for 1h in a cell incubator. Place 200 µl of TMB color developing solution and 5 µl of small chamber culture medium in a 96-well plate and let stand for 30 min. The OD value of each 96-well sample was measured using a microplate reader to calculate the HRP content of the lower chamber. The amount of HRP bleed is expressed in pmol per cubic centimeter of HRP per hour (pmol/cm2/h).

### Western Blot Assays

The total protein of the cells was extracted with RIPA buffer (Beyotime Institute of Biotechnology, Jiangsu Province) supplemented with protease inhibitors (10 mg/mL aprotinin, 10 mg/mL PMSF and and 50 mM sodium orthovanadate).The protein concentration was then determined using the BCA protein assay kit (Jiangsu Beiyang Institute of Biotechnology, China).The same amount of protein (40g) was loaded for SDS-PAGE electrophoresis and then transferred to Millipore (Shanghai, China) and sealed in Tris buffer/Tween 20 (TBST) containing 5% skimmed milk powder for 2 hours at room temperature.Incubate primary antibody overnight at 4°C.The membrane was washed three times with TBST and then incubated at room temperature with conjugated HRP secondary antibodies for 2 hours.ECL (enhanced chemiluminescence kit, Santa Cruz Biotechnology, Dallas, TX) detection system (Thermo Scientific, Beijing, China) was used for detection. Scan with Chemi Imager 5500 V2.03. GAPDH was used as internal reference to determine the expression level of the target protein. Antibody used are provided in Table S3.

### Immunofluorescence Assays

Cell slides were fixed with paraformaldehyde in the dark for 30 min and washed three times with PBST. After penetrating the membrane with Trixton-100 for 10 min, the slides were washed with PBST. After blocking with 5% BSA for 2 h, it was incubated with the corresponding antibodies (1:50; Life Technologies) of ZO-1, occludin and claudin-5 overnight. After reheating at room temperature for half an hour, wash the primary antibody with PBST, and then apply its corresponding secondary antibody (1:500; Beyotime Institute of Biotechnology, Jiangsu, China) at room temperature for 2h. Wash three times with PBST for 10 min each time. The nuclei were stained with DAPI for 5 min. After the staining was completed, PBST was washed three times, and the slides were sealed with 50% glycerol. Observe and take pictures under a confocal microscope.

### Methylated RNA immunoprecipitation sequencing (MeRIP-seq) and MeRIP-qPCR

We used RiboMinus™ Eukaryote Kit v2 (A15020, Invitrogen) to extract and purify 50 pounds of total RNA to extract ribosomal RNA from total RNA. Next, RNA lysis reagent (AM8740, Invitrogen) was used to cut RNA into approximately 100-nt fragments. Approximately 1/10 of the fragmented RNA was saved as an input control for further RNA sequencing by KangChen (Shanghai, China). The remaining cells were incubated with anti-m^6^A antibody (ab208577, Abcam) at 4°C for 1 hour, and then mixed with pre-washed Pierce™ Protein A/G magnetic beads (88,803, Thermo Scientific) in 4°C immunoprecipitation buffer overnight. The m^6^A antibody was digested with proteinase K digestion buffer, and the methylated RNA was purified and further sequenced and qPCR by KangChen (Shanghai, China).

### m^6^A dot blot assay

The Poly (A) + RNA was first denatured by heating at 65°C for 5 minutes and then transferred to A cellulose nitrate membrane (Amersham, GE Healthcare, US) using A Bio-dot device (US Bio-RAD).The membranes were then uV-crosslinked, sealed, and incubated overnight at 4°C with m^6^A antibody (1:1000, Abcam, USA), and then incubated with HRP-conjugated goat anti-mouse IgG (1:300, Proteintech, USA).Finally, membranes can be visualized with chemiluminescence systems (Bio-RAD, US).The membrane was stained with 0.02% methylene blue (MB) in 0.3 M sodium acetate (pH 5.2) to ensure consistency across groups.

### M^6^A mutation assays

The online tool SRAMP (http://www.cuilab.cn/sramp/) was used to predict possible m^6^A sites. The full-length transcript of CPEB2, CPEB2 CDS region, CPEB2 3’-UTR three’ untranslated region, and m^6^A motif deleted CDS or 3’-UTR region were cloned into pcDNA3.1 for RNA pull down experiment. The specific sequences were shown in Table S5.

### RNA immunoprecipitation (RIP) assay

Whole cell lysates from different groups were collected and incubated with magnetic bead RIP buffer containing anti-human argonaute 2 (Ago2) antibody (Millipore, Billerica, MA, USA) and anti-human CPEB2 antibody (Proteintech) overnight. The negative control group was incubated with normal mouse IgG (Millipore). Samples were incubated with proteinase K buffer to isolate immunoprecipitated RNA. The RNA concentration was measured using a spectrophotometer (NanoDrop, Thermo Scientific), and the RNA quality was evaluated using a bioanalyzer (Agilent, Santa Clara, CA, USA). Then purified and reverse transcribed the RNA, detect RNA enrichment by qRT-PCR.

### RNA pull-down assay

The interaction between CPEB2 and SRSF5 was tested by Pierce Magnetic Magnetic protein pull-down kit (Thermo Fisher) according to the manufacturer’s instructions. Biotin-labeled SRSF5 (Bio-SRSF5) was transcribed *in vitro* with biotin-RNA-labeled mixture (Roche) and T7 RNA polymerase (Promega). Bio-SRSF5 and Bio-antisense RNA (NC) were incubated with GEC lysates and then washed. Magnetic beads were added to prepare probe-magnetic bead complexes.The recovered protein was analyzed by western blot with GAPDH as the control.

### Nascent RNA Capture

Nascent RNAs was detected by Click-iT nascent RNA capture kit (Thermo Fisher Scientifific, USA). 5-ethymyl uridine (EU) was clicked into nascent RNAs, then streptavidin magnetic beads captured EU-nascent RNAs for the following quantitative real-time PCR.

### RNA stability measurement

Cells were cultured in medium containing 5 µg/ml actinomycin D (Act D, NobleRyder, China).Then extracted total RNA at different time points, and detected by real-time fluorescence quantitative PCR. Compared to zero time, the half-life of RNA is determined by its level being reduced to 50% at a certain point.

### Human mRNA PCR Arrays

For mRNA PCR arrays analysis, sample preparation and array hybridization were performed by KangChen Bio-tech (Shanghai, China).

### Minigene assay

The ETS1 exon 7 minigene was constructed by insertion of exons and its flanking intron region into pGint Vector. ETS1 exon 7 and its flanking intron region were amplified with ETS1 minigene-F and minigene-R from genome DNA and inserted into the intron. The specific primers of complete EGFP RNA were identified to determine the splicing efficiency by qPCR. Primers are shown in Table S1.

### Reporter Vector Construction and Dual Luciferase Reporter Assays

Reporter vector construction and dual luciferase reporter assays were conducted as previously reported^[56]^. The potential binding sequence and corresponding mutant sequence of ZO-1 3’-UTR, occludin 3’-UTR and Claudin-5 3’-UTR of P51-ETS1 were amplified by PCR, and cloned into the targeted expression vector of pmirGLO double luciferase miRNA (Promega, Madison, Wisconsin, USA) to construct the wild-type and mutant luciferase reporter vectors (Generay Biotech Co., Shanghai, China). Lipofectamine 3000 was used to co-transfect HEK-293T cells with wild-type and mutant luciferase reporter vectors as well as P51-ETS1-NC or P51-ETS1.Relative luciferase activity was expressed as the ratio of firefly luciferase activity to sea kidney luciferase activity using a dual luciferase reporting kit (Promega, Madison, Wisconsin, USA) after 48 h.

### ChIP Assays

The ChIP assays were performed using the Simple Chip Enzymatic Chromatin IP kit (Cell Signaling Technology, Danvers, MA, USA) according to the manufacturer’s instructions.GECs were crosslinked with EBM-2 containing 1% formaldehyde for 10 minutes, and incubated with glycine at room temperature for 5 minutes to stop the reaction crosslinking. Add PMSF containing lysate to the cells in the dish to fully lyse. The chromatin in the lysed cells is digested with nuclease. The immunoprecipitation was incubated with 2 µg P51-ETS1 antibody (Santa, USA). Protein G agarose beads were subjected to a cross-linking reaction with their DNA while 2% of the lysate was used as a positive control Input. Add 5 mol/L NaCl and proteinase K to the DNA cross-linking solution to purify the DNA. The purified DNA was used for PCR amplification to verify the binding of P51-ETS1 to the promoter regions of ZO-1, occludin, and claudin-5. Primers used for ChIP PCR are shown in Table S4.

### Evaluation of Apoptosis by Flow Cytometry

After establishing the BTB model *in vitro*, 10 mDOX was added to the upper chamber of Transwell (Beyotime Institute of Biotechnology, Jiangsu, China).The apoptosis rate of U251 cells inoculated in the lower compartment was measured using a membrane-bound V-PE/7AAD kit (Southern Biotech, Birmingham, Ala, USA) after 12h.Then washing with phosphate buffer and centrifugation twice, U251 cells in the lower chamber were resuspended in the membrane-bound V binding buffer.The resuspended cells were stained with membrane-bound V-PE / 7AAD at room temperature for 15 minutes in darkness.Cell samples were obtained by FACScan (BD Biosciences) for apoptotic rate and analyzed by Cell Quest 3.0 software.

### Statistical Analysis

GraphPad Prism v5.01 software was used for statistical analysis. Statistical analysis of data was performed using the Student’s t test in significant differences between two groups. using one-way ANOVA followed by Dunnett’s post hoc test for three or more groups. Data were presented as the mean ± SD. *P* < 0.05 was considered as statistically significant.

## Results

### 1. METTL3 is highly expressed in GECs, knockdown METTL3 increases BTB permeability

In this study, qRT-PCR was used to detect METTL3, METTL14, WTAP, and METTL4 mRNA expression levels in an *in vitro* BTB model. Compared with the AECs group, METTL3 expression levels in GECs were significantly increased (Figure S1a). It was found that METTL3 was highly expressed in glioblastoma compared with normal samples in TCGA database which was analyzed by UALCAN (http://ualcan.path.uab.edu) (Figure S1b); The clinical samples with high expression of METTL3 in glioblastoma had a lower survival rate than those with low expression of METTL3 (Figure S1c), suggesting that the increased expression of METTL3 is associated with the poor prognosis of malignant glioma. In this study, it was found that in the *in vitro* BTB model the METTL3 mRNA and protein expression levels in the GECs group were significantly higher than those in the AECs group (Figure 1a, b). The TEER value and horseradish peroxidase (HRP) flux were measured after silencing METTL3 which was in GECs to analyze the integrity and permeability of BTB. The results showed that compared with sh-METTL3-NC group, TEER value of sh-METTL3 group decreased significantly and the HRP flux increased significantly (Figure 1c, d). Detection of changes in the expression levels of tight junction-related proteins ZO-1, occludin, and claudin-5 showed that compared with the sh-METTL3-NC group, the expression levels of ZO-1, occludin, and claudin-5 in the sh-METTL3 group were significant lower (Figure 1e,f). The results of immunofluorescence staining are shown in Figure 1g. ZO-1, occludin, claudin-5 in the control group and sh-METTL3-NC group showed a continuous distribution at the tight junctions of GECs, in the sh-METTL3 group, no continuity distribution of the above proteins.

**Fig. 1.**
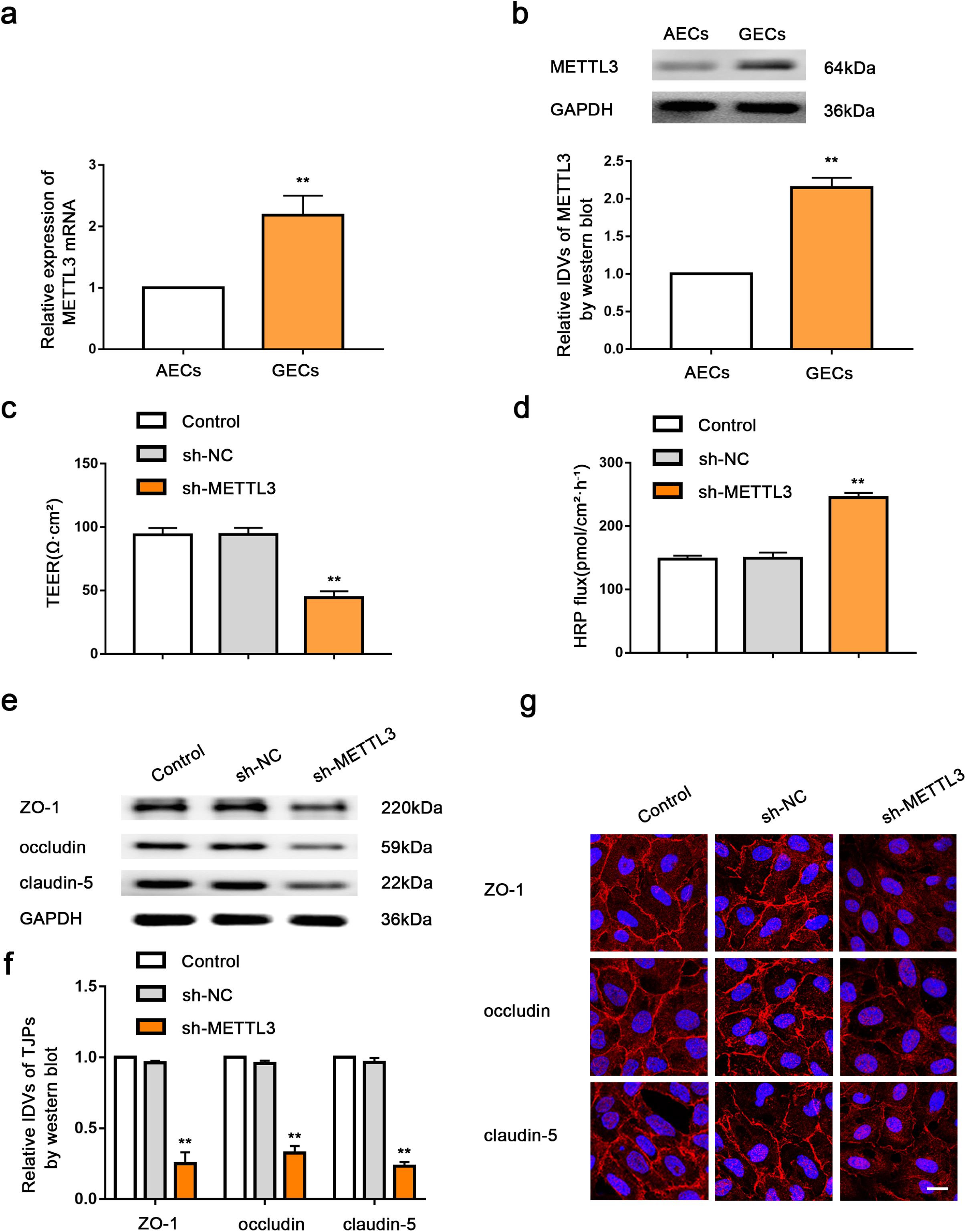
Knockdown of METTL3 increased BTB permeability *in vitro*. **a** Relative mRNA levels of METTL3 in AECs (the NHAs co-cultured ECs) and GECs (the glioma cells co-cultured ECs) were determined by qRT-PCR. **b** Relative protein levels of METTL3 in AECs and GECs were determined by western blot. Data represented as mean ± SD (n = 3). \*\**P* < 0.01 vs. AECs group. **c, d** The permeability and integrity of the METTL3 knockdown BTB model *in vitro* were detected by TEER values and HRP flux.e The expressions of ZO-1, occludin, and claudin-5 in the METTL3 knockdown GECs detected by western blot. Data represented as mean ± SD (n = 3). \*\**P* < 0.01 vs. sh-METTL3-NC group. **f** The distributions of ZO-1, occludin and claudin-5 in the METTL3 knockdown GECs were observed by immunofluorescent staining. Scale bar represents 50 µm.

### 2. METTL3 mediated m^6^A methylation of CPEB2 mRNA

To identify the molecular mechanism by which METTL3 inhibits BTB permeability, we conducted MeRIP-seq in GECs with stable METTL3 downregulation, used sh-NC as controls. The results showed that sh-METTL3 group had 8153 m^6^A peaks, which was 1125 peaks less than sh-NC (9278 m^6^A peaks), (Figure 2a). We analyzed the top 1000 most significant peaks and found that there was a consensus sequence RRACH (Figure 2b), and the m^6^A enrichment region of CPEB2 located around the stop codon (Figure 2c). Differential genes were analyzed by DAVID (https://david.ncifcrf.gov/home.jsp) for signal pathway analysis (Figure S2a) and GO (Gene Ontology) analysis (Figure S2b). Through MeRIP-seq data analysis, 10 related genes were finally obtained (Figure 2d). After silencing METTL3, the mRNA expression levels of these 10 genes were detected. Compared with the sh-NC group, CPEB2 mRNA expression levels were significantly reduced, (Figure S2c). According to MeRIP-seq data, methylation sites of CPEB2 were found. In order to determine whether m^6^A modification of CPEB2 is mediated by METTL3, we first measured the overall m^6^A level in the control group and sh-METTL3 group by m^6^A dot blot assay, and found that the overall m^6^A level in the sh-METTL3 group was significantly reduced(Figure 2d). In addition, m^6^A enrichment in CPEB2 was determined by MeRIP-qPCR. Compared with IgG control, m^6^A specific antibody could stably enrich CPEB2 mRNA. In addition, we found that the amount of CPEB2 modified by m^6^A after silencing METTL3 was significantly reduced(Figure 2e). Therefore, we believe that METTL3 can affect the overall level of m^6^A, especially the level of CPEB2 m^6^A.

**Fig. 2.**
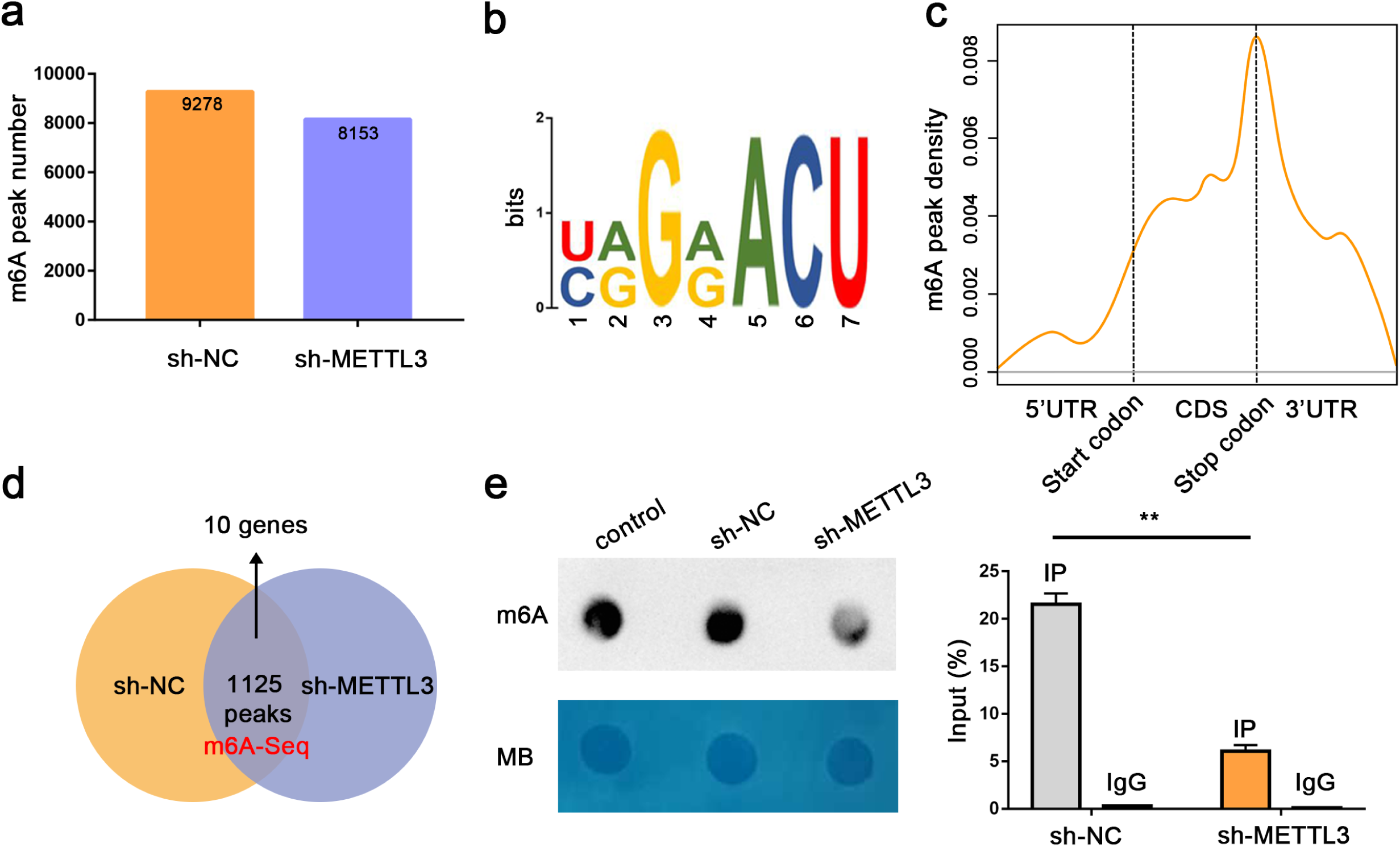
m6A-Seq profiling of GECs and identified CPEB2 as a target of METTL3-mediated m^6^A modification. **a** m^6^A peak numbers in sh-METTL3 GECs vs sh-NC group. **b** Top consensus motif identified in GECs. **c** The normalized distribution of m^6^A peaks across the start codon, CDS, stop codon of mRNAs for m^6^A peaks. **d** Venn diagram of m^6^A-seq. **e** m^6^A dot blot assay of METTL3 silenced GECs. Methylene blue staining served as a loading control. **f** Relative enrichment of CPEB2 m^6^A modification in the METTL3 knockdown GECs vs. Sh-NC group were analyzed using qRT-PCR.

### 3. CPEB2 is highly expressed in GECs, silencing CPEB2 increases BTB permeability; IGF2BP3 combined with CPEB2 mRNA 3’-UTR increases its mRNA stability

In the *in vitro* BTB model, the expression of CPEB2 mRNA and protein in the GECs group was significantly higher than that in the AECs group (Figure 3a-c). The results of detecting TEER value and HRP flux after silencing CPEB2 showed that compared with the sh-CPEB2-NC group, the TEER value of the sh-CPEB2 group was significantly reduced and the HRP flux was significantly increased (Figure 3d, e). The results of the effect of sh-CPEB2 on the expression of tight junction-related proteins ZO-1, occludin and claudin-5 showed that the expression of ZO-1, occludin and claudin-5 in the sh-CPEB2 group was significantly reduced compared with that of the sh-CPEB2-NC group (Figure 3f). Immunofluorescence staining results showed that ZO-1, occludin and Claudin-5 of the control group and the sh-CPEB2-NC group were continuously distributed in the GECs tight junction, as shown in Figure 3g, while in the sh-CPEB2 group, the above proteins were not continuously distributed (Figure 3h).

**Fig. 3.**
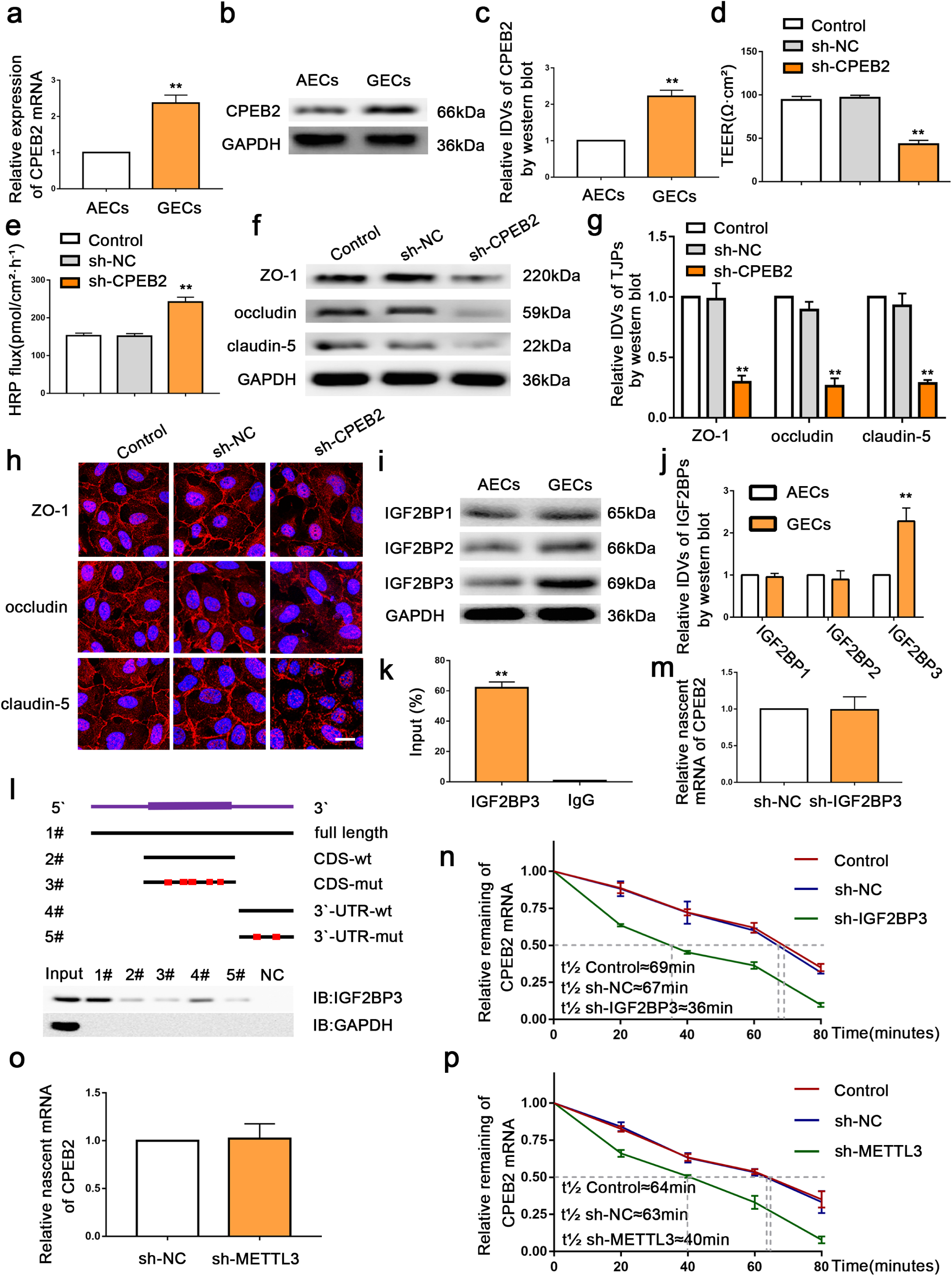
Knockdown of CPEB2 increased BTB permeability *in vitro*. **a** Relative mRNA levels of CPEB2 in AECs and GECs were determined by qRT-PCR. **b,c** Relative protein levels of CPEB2 in AECs and GECs were determined by western blot. Data represented as mean ± SD (n = 3). \*\**P* < 0.01 vs. AECs group. **d,e** The permeability and integrity of the CPEB2 knockdown BTB model *in vitro* were detected by TEER values and HRP flux. **f,g** The expressions of ZO-1, occludin, and claudin-5 in the CPEB2 knockdown GECs detected by western blot. Data represented as mean ± SD (n = 3). \*\**P* < 0.01 vs. sh-CPEB2-NC group. h The distributions of ZO-1, occludin, and claudin-5 in the CPEB2 knockdown GECs were observed by immunofluorescent staining. Scale bar represents 50 µm. **i,j** Relative protein levels of IGF2BP1, IGF2BP2, IGF2BP3 in AECs and GECs were determined by western blot. Data represented as mean ± SD (n = 3). \*\**P* < 0.01 vs. AECs group. k RNA immunoprecipitation assay was performed with normal mouse IgG or anti-IGF2BP3 antibody in GECs. Relative enrichment of CPEB2 was determined by qRT-PCR. l Immunoblotting of IGF2BP3 with cell lysate (Ly.), full-length biotinylated-CPEB2 (#1), the CPEB2 CDS region with or without m^6^A motif mutation (#2, #3), the CPEB2 3’-UTR region with or without m^6^A motif mutation (#4, #5), and beads only (NC) in CPEB2 cells. m Relative expression levels of nascent CPEB2 in the IGF2BP3 knockdown GECs were detected using qRT-PCR. **n** Relative expression levels of CPEB2 in the IGF2BP3 knockdown GECs treated with actinomycin D at different time points were analyzed using qRT-PCR. **o** Relative expression levels of nascent CPEB2 in the METTL3 knockdown GECs were detected using qRT-PCR. **p** Relative expression levels of CPEB2 in the METTL3 knockdown GECs treated with actinomycin D at different time points were analyzed using qRT-PCR.

This study further investigated the expression of IGF2BP1, IGF2BP2 and IGF2BP3 in AECs and GECs in the m^6^A binding protein IGF2BP family, and found that the expression level of IGF2BP3 in GECs was significantly increased (Figure 3i,j). After IGF2BP3 was silenced, the TEER value of the sh-IGF2BP3-NC group was significantly reduced, HRP flux was significantly increased, and the expression levels and continuous distribution of tight junction related proteins ZO-1, occludin, and claudin-5 were significantly reduced (Figure S3). The results of RIP and RNA pulldown experiments showed that IGF2BP3 directly binds to the 3’-UTR end of CPEB2 mRNA (Figure k,l). The results of the nascent RNA experiment are shown in Figure 3m. Compared with sh-IGF2BP3-NC, there was no significant change in the CPEB2 nascent RNA in the sh-IGF2BP3 group. After the addition of actinomycin D, the half-life of CPEB2 mRNA was measured at different times. Compared with the sh-IGF2BP3-NC group, the half-life of CPEB2 mRNA in the sh-IGF2BP3 group was significantly shortened (Figure 3n). After silencing METTL3, the nascent RNA experiment was performed, and the results showed that there was no statistically significant change in CPEB2 nascent RNA in the sh-METTL3 group compared with sh-NC (Figure 3o); the results of the half-life showed that compared with the sh-NC group, the sh-CPEB2 mRNA half-life of the METTL3 group was significantly shortened (Figure 3p).

### 4. SRSF5 is highly expressed in GECs, and silencing SRSF5 increases the permeability of BTB; CPEB2 enhances the stability of SRSF5 mRNA and regulates the permeability of BTB

In this study, mRNA PCR array analysis was performed after METTL3 was silenced in GECs and verified by qRT-PCR, results showed that the expression level of SRSF5 was significantly decreased (Figure S4a). In the *in vitro* BTB model, the expression of SRSF5 mRNA and protein in the GECs group was significantly higher than that in the AECs group (Figure 4a-c). Compared with the sh-NC group, the TEER value of the sh-SRSF5 group was significantly reduced, the HRP flux was significantly increased, and the protein expression levels of ZO-1, occludin, and claudin-5 were significantly reduced (Figure 4d-g). Immunofluorescence staining showed that in the sh-SRSF5 group, the above proteins showed a discontinuous distribution (Figure 4h). RIP and RNA pulldown assays showed that CPEB2 binds to SRSF5 mRNA (Figure 4i, j). Compared with the sh-CPEB2-NC group, there was no significant change in nascent RNA (Figure 4k), and the SRSF5 mRNA half-life of the sh-CPEB2 group was significantly shortened (Figure 4l). In order to determine that CPEB2-mediated SRSF5 regulates BTB permeability through m^6^A modification, this study mutated the CPEB2 methylation site and conducted a rescue experiment. The results are shown in Figure S4b-f. Compared with CPEB2-WT, the expression levels of SRSF5, ZO-1, occludin, and claudin-5 in the CPEB2-m^6^A-mut group were significantly reduced. Overexpression of CPEB2 after the m^6^A mutation was found. Compared with the sh-NC group, the expression levels of the above proteins were not significant. Variety. TEER value, HRP flux and immunofluorescence results showed that the BTB permeability was significantly increased after mutating the m^6^A site. On this basis, the overexpression of CPEB2 showed no significant change in the permeability of BTB compared with sh-NC group.

**Fig. 4.**
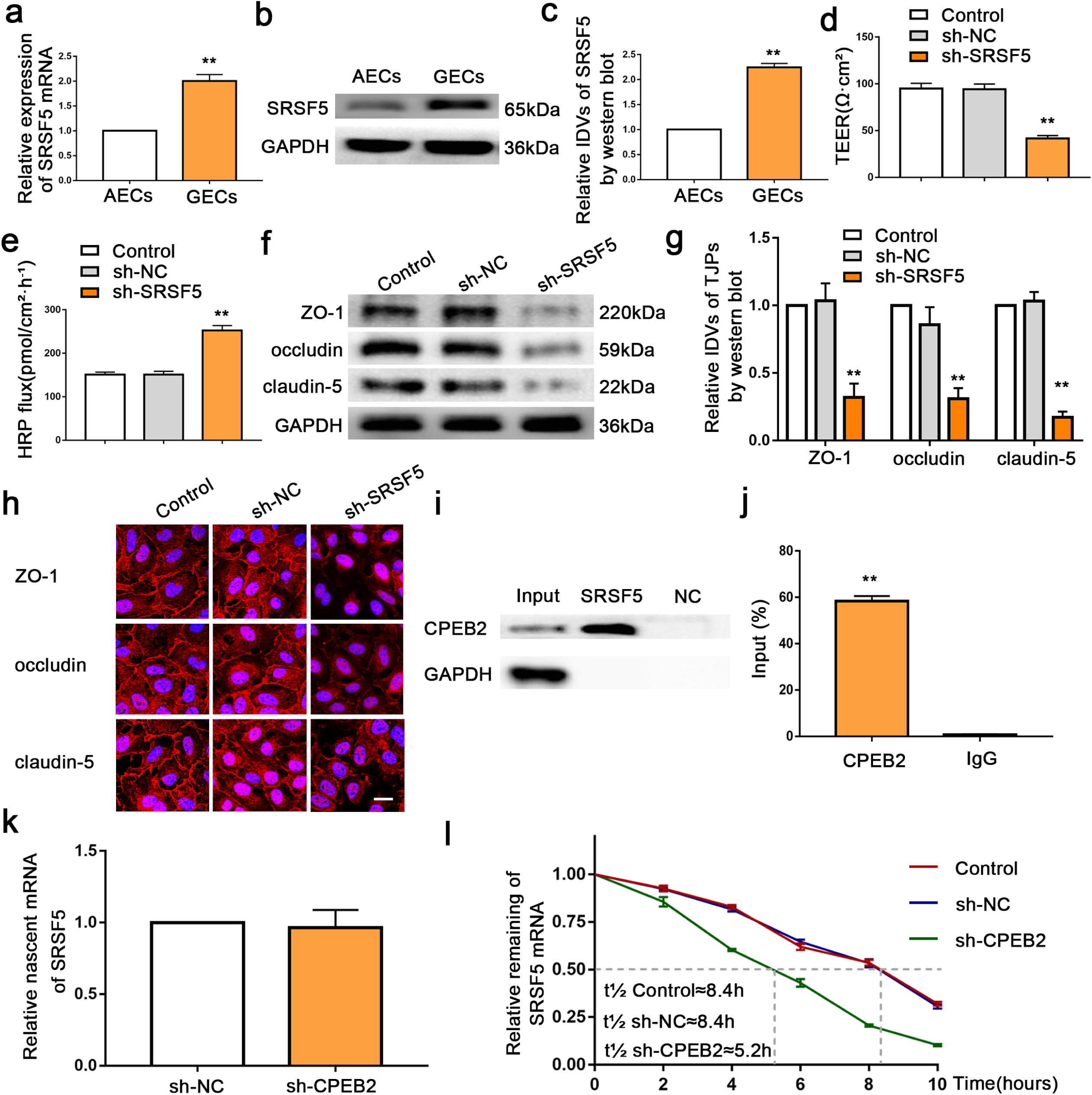
Knockdown of SRSF5 increased BTB permeability *in vitro*. **a** Relative mRNA levels of SRSF5 in AECs and GECs were determined by qRT-PCR. **b,c** Relative protein levels of SRSF5 in AECs and GECs were determined by western blot. Data represented as mean ± SD (n = 3). \*\**P* < 0.01 vs. AECs group. **d,e** The permeability and integrity of the SRSF5 knockdown BTB model *in vitro* were detected by TEER values and HRP flux. **f,g** The expressions of ZO-1,occludin and claudin-5 in the SRSF5 knockdown GECs detected by western blot. Data represented as mean ± SD (n = 3). \*\**P* < 0.01 vs. sh-SRSF5-NC group. **h** The distributions of ZO-1, occludin and claudin-5 in the SRSF5 knockdown GECs were observed by immunofluorescent staining. Scale bar represents 50 µm. **i** RNA immunoprecipitation assay was performed with normal mouse IgG or anti-CPEB2 antibody in GECs. **j** Relative enrichment of SRSF5 was determined by qRT-PCR. Data represented as mean ± SD (n = 3). \*\**P* < 0.01 vs. anti-IgG group. **k** Relative expression levels of nascent SRSF5 in the CPEB2 knockdown GECs were detected using qRT-PCR. **l** Relative expression levels of SRSF5 in the CPEB2 knockdown GECs treated with actinomycin D at different time points were analyzed using qRT-PCR.

### 5. P51-ETS1 is highly expressed in GECs, silencing P51-ETS1 increases the permeability of BTB

ETS1 contains multiple splicing variants. In this study, two siRNAs (S1, S2) were designed in the VII exon of ETS1, and siRNAs (S3, S4) were designed in the VI exon and the VIII exon respectively (Figure S5a), qRT-PCR and Western blot detection results show that compared with the corresponding si-NC group, si-S1 and si-S2 significantly reduce the mRNA and protein expression of full-length P51-ETS1, while truncated P42 -ETS1 mRNA and protein expression levels increased significantly; si-S3 and si-S4 significantly decreased the mRNA and protein expression of P51-ETS1 and P42-ETS1 (Figure S5b, c), suggesting that P51-ETS1 and P42-ETS1 differ by Exon VII. In the *in vitro* BTB model, the expression levels of P51-ETS1 mRNA and protein in the GECs group were significantly higher than those in the AECs group, and the expression of P42-ETS1 mRNA and protein were significantly lower than in the AECs group (Figure 5a-c). Compared with the sh-NC group, the TEER value of the sh-P51-ETS1 group decreased significantly and the HRP flux increased significantly; the pre-P51-ETS1 group had a significantly increased TEER value and significantly decreased HRP flux (Figure 5d, e). The effect of silence and overexpression of P51-ETS1 on the expression of tight junction-related proteins ZO-1, occludin, claudin-5 is shown in Figure 5f,g, compared with the sh-NC group, the expression of ZO-1, occluding, claudin-5 in sh-P51-ETS1 group was significantly reduced, and the expression of ZO-1, occludin and claudin-5 in the pre-P51-ETS1 group was significantly increased. Immunofluorescence staining results showed that ZO-1, occludin, claudin-5 in the control group and sh-P51-ETS1-NC group showed a continuous distribution at the tight junctions in GECs, in the sh-P51-ETS1 group, the continuous distribution of the above-mentioned proteins was not seen; while in the pre-P51-ETS1 group, the distribution of the above-mentioned proteins was more closely connected (Figure 5h). We established GECs stably expressing sh- P51-ETS1+pre-P51-ETS1, results showed that TEER value, HRP flux and ZO-1, occludin, claudin-5 expression levels were no significant change compared with the sh-NC group; while P51-ETS1+pre-P42-ETS1 group, TEER value was significantly reduced, HRP permeability was significantly increased, ZO-1, occludin, claudin-5 expression levels were significantly reduced (Figure S5d-g). Immunofluorescence staining results showed that sh-P51-ETS1+pre-P51-ETS1 group permeability did not change significantly compared with the sh-NC group; P51-ETS1+pre-P42-ETS1 group, permeability was significantly reduced compared to the sh-NC group (Figure S5h).

**Fig. 5.**
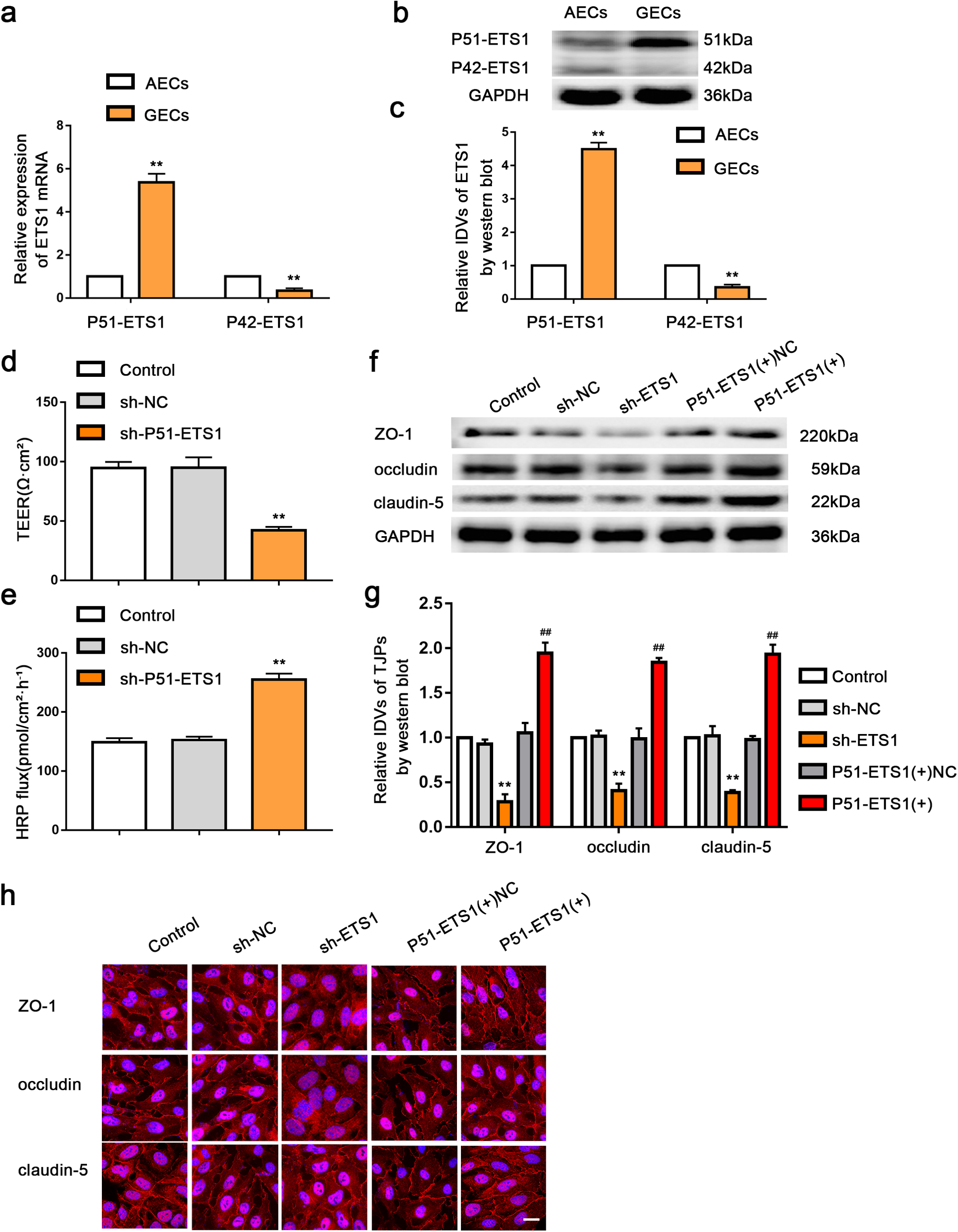
Knockdown of P51-ETS1 increased BTB permeability *in vitro*. **a** Relative mRNA levels of P51-ETS1 and P42-ETS1 in AECs and GECs were determined by qRT-PCR. **b,c** Relative protein levels of P51-ETS1 and P42-ETS1 in AECs and GECs were determined by western blot. Data represented as mean ± SD (n = 3). \*\**P* < 0.01 vs. AECs group. **d,e** The permeability and integrity of the P51-ETS1 knockdown BTB model *in vitro* were detected by TEER values HRP flux. **f,g** Effects of ETS1 expression changes on the ZO-1, occludin and claudin-5 protein levels were detected by Western blot. Data represented as mean ± SD (n = 3). \*\**P* < 0.01 vs. sh-NC group, ##*P* < 0.01 vs. pre-NC group. **h** Effects of ETS1 expression changes on the distributions of ZO-1, occludin and claudin-5 were detected by immunofluorescent staining. Scale bar represents 50 µm.

### 6. SRSF5 induces ETS1 exon VII inclusion, and P51-ETS1 binds to the TJPs promoter region

In this study, we used ESEfinder (http://exon.cshl.edu/ESE/) to predict the binding motif of SRSF5 and ETS1 mRNA, and found this motif (CCACAAG) on exon 7 of ETS1 mRNA (Figure S6a). In order to determine whether SRSF5 specifically binds to this motif to regulate AS, RIP assays revealed that SRSF5 can directly bind to ETS1 mRNA (Figure S6b). The study further determined whether ETS1 was the AS target of SRSF5. Compared with the sh-SRSF5-NC group, the expression levels of P51-ETS1 mRNA and protein in the sh-SRSF5 group were significantly reduced, and P42-ETS1 mRNA and protein were significantly increased (Figure 6a-c). we established GECs stably expressing sh-SRSF5+pre-P51-ETS1, TEER value, HRP flux and the expression levels of ZO-1, occludin, and claudin-5 were not significantly changed compared with the sh-NC group; while in sh-SRSF5+pre-P42-ETS1 group, TEER value was significantly reduced, HRP permeability was significantly increased, the expression levels of ZO-1, occludin and claudin-5 were significantly lower than those of the sh-NC group (Figure S6c-f). The immunofluorescence results showed that BTB permeability did not change significantly in sh-SRSF5+pre-P51-ETS1 group compared with the sh-NC group; in sh-SRSF5+pre-P42-ETS1 group, the BTB permeability was significantly reduced compared to the sh-NC group (Fig. S6g). In order to clarify whether SRSF5 can induces ETS1 exon VII inclusion, we constructed the ETS1 exon7 small gene reporting system, after skipping P42-ETS1 exon7 (ETS1 ex7-), EGFP retained the entire ORF, while P51-ETS1 containing exon7 destroyed ORF and reduced EGFP expression (Figure S6h). As shown in the figure, silencing SRSF5 caused a significant increase in the EGFP signal (Figure S6i), indicating that SRSF5 promoted the retention of exon VII of ETS1. Primers that specifically recognize intact EGFP RNA were used to assess splicing efficiency; qRT-PCR showed that silencing SRSF5 significantly increased ETS1 splicing efficiency (Figure S6j). Dual-luciferase reporter gene assays showed the activities of the ZO-1, occludin, and claudin-5 promoters were significantly reduced in the pEX3-P51-ETS1 group, compared with those in the pEX3 empty vector group (Figure 6d-f). And the chromatin immunoprecipitation (ChIP) assays proved P51-ETS1 directly bind to the ZO-1, occludin and claudin-5 promoter region binding sites, did not interact with the control region (Figure 6g-i). Then we established sh-CPEB2+sh-SRSF5 GECs, compared with the sh-NC group, TEER value was significantly reduced, HRP permeability was significantly increased, the expression level of P51-ETS1 was significantly decreased, the expression level of P42-ETS1 was significantly increased, and the expression levels of ZO-1, Occludin and Claudin-5 were significantly decreased.; while in sh-CPEB2+pre-SRSF5 group, there was no significant change in the above protein expression levels compared with the sh-NC group (Figure S6k-n). The immunofluorescence results showed that CPEB2 and SRSF5 double-silence significantly reduced BTB permeability compared with the sh-NC group; as for sh-CPEB2+pre-SRSF5 group, there was no significant change in BTB permeability, compared with the sh-NC group (Figure S6o).

**Fig. 6.**
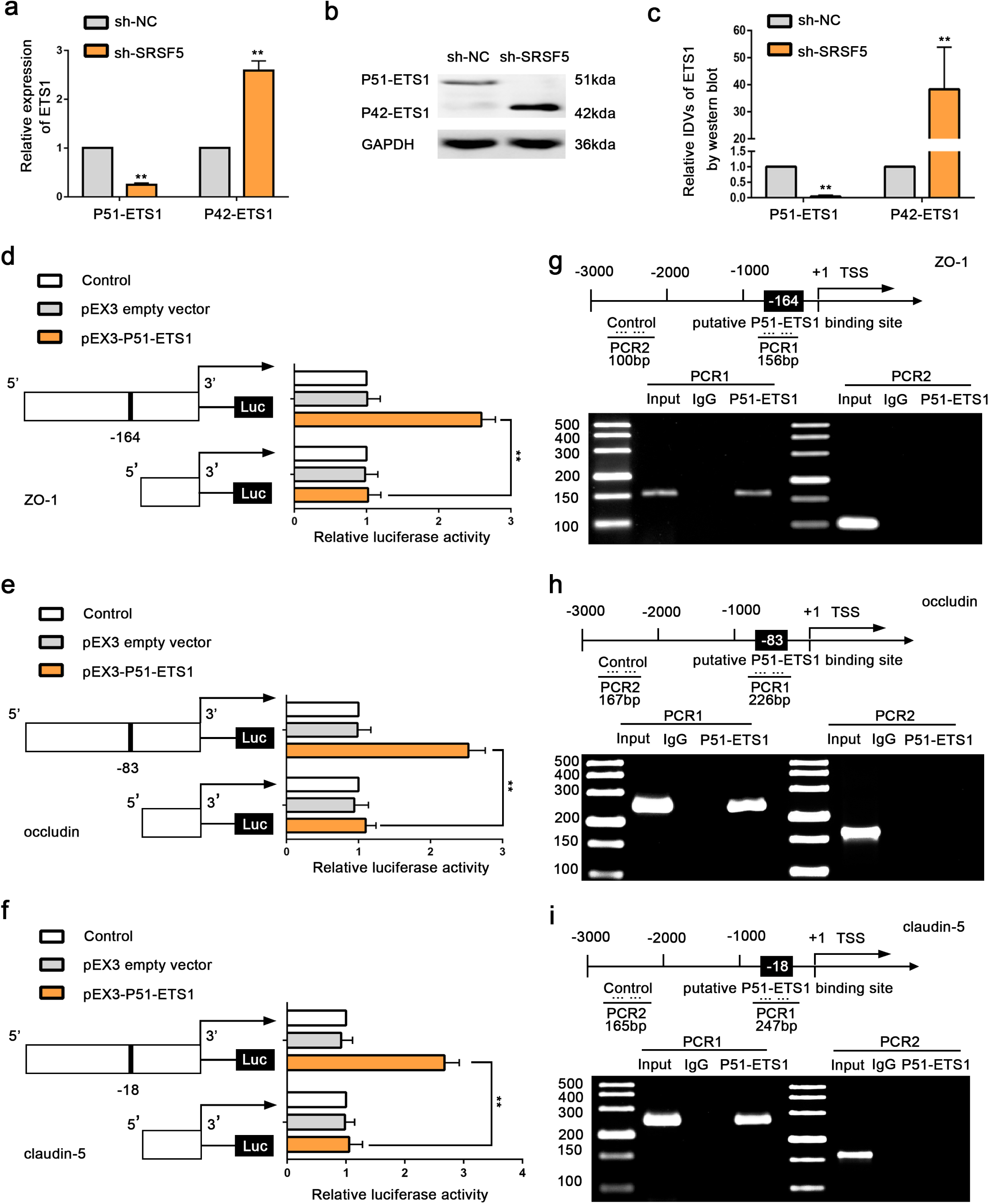
SRSF5 inhibits ETS1 AS events and P51-ETS1 binds to the TJPs promoter region. **a** Relative mRNA levels of P51-ETS1 and P42-ETS1 in GECs and sh-SRSF5 GECs were determined by qRT-PCR. **b-c** Relative protein levels of P51-ETS1 and P42-ETS1 in GECs and sh-SRSF5 GECs were determined by western blot. Data represented as mean ± SD (n = 3). \*\**P* < 0.01 vs. GECs group. **d-f** Dual luciferase reporter assays were performed to determine the binding sites of ETS1 and ZO-1, occludin and claudin-5 in HEK293T cells. **g-i** ZO-1, occludin and claudin-5 promoter region 3,000 bp upstream of the transcription start sites (TSSs), which were designated as +1.PCR1 represents the unbound negative control group, and PCR2 represents the binding site of P51-ETS1 to the tightly bound proteins.

### 7. The combined application of multiple targets promotes doxorubicin (Dox) to induce glioma cell apoptosis

In this study, the effects of silencing METTL3, IGF2BP3, CPEB2, SRSF5 and P51-ETS1 alone or combined on apoptosis of U251 glioma cells were investigated. The results are shown in Figure 7a, b, compared with the Dox group, the Dox-treated silenced CPEB2, SRSF5, and ETS1, and the CPEB2, SRSF5, P51-ETS1 triple-silenced group had a significant increased apoptosis rate of U251 glioma cells; when CPEB2, SRSF5 and P51-ETS1 were used in combination, the effect of inducing apoptosis of glioma cells was more statistically significant than that when applied alone. Figure 7c is a schematic diagram of the mechanism that CPEB2 m^6^A modification affects the stability of the splicing factor SRSF5 and regulates the permeability of BTB.

**Fig. 7.**
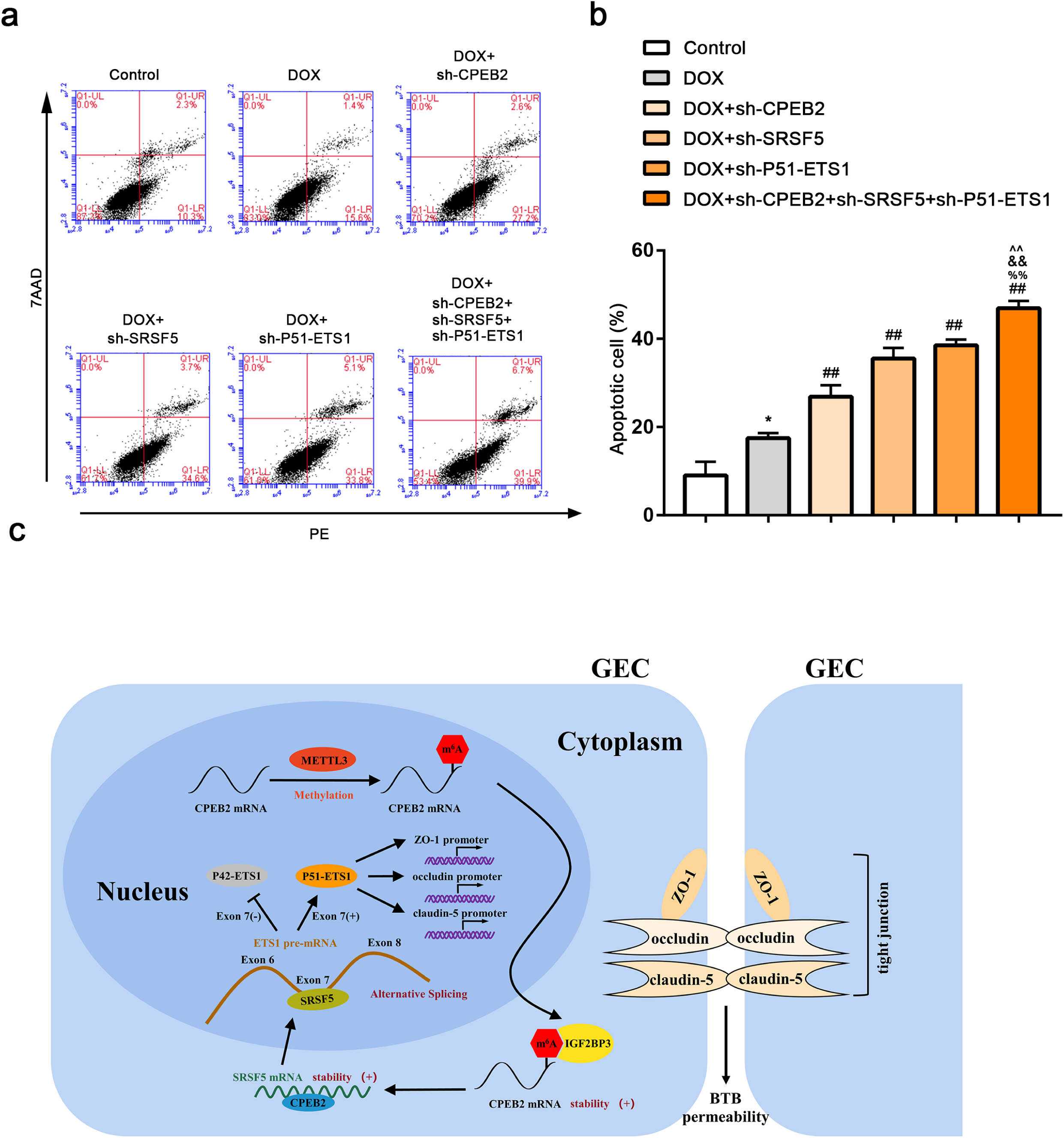
The apoptosis changes of U251 cells induced by combined treatment of METTL3, IGF2BP3, CPEB2, SRSF5, and P51-ETS1 with doxorubicin. **a** Representative flow cytometric detection a and analysis of the U251 cell apoptosis rates in different groups. **b** Data represented as mean ± SD (n = 3). \**P*<0.05 vs. control group, ##*P* < 0.01 vs. DOX group, %% *P* < 0.01 vs. DOX+ sh-CPEB2 group, &&*P* < 0.01 vs. sh-SRSF5 group, ^^*P* < 0.01 vs. sh-P51-ETS1 group. **c** The schematic diagram of the mechanism that CPEB2 m^6^A modification affects the stability of the splicing factor SRSF5 and regulates the permeability of BTB.

## Discussion

Because of the presence of the blood tumor barrier (BTB), it is difficult for large-molecule chemotherapeutic drugs to reach the glioma tissue to exert therapeutic effects. Selective opening of BTB is an effective strategy to improve the chemical efficacy of glioma. This study found that METTL3, IGF2BP3, CPEB2, SRSF5, and P51-ETS1 were significantly up-regulated in GECs. Silencing METTL3, IGF2BP3, CPEB2, SRSF5, and P51-ETS1 significantly reduced the expression levels of ZO-1, occludin, and claudin-5, and increased the permeability of BTB. METTL3 promotes CPEB2 mRNA m^6^A methylation, IGF2BP3 enhances CPEB2 mRNA stability by recognizing m^6^A methylation sites, CPEB2 enhances SRSF5 mRNA stability and regulates SRSF5-mediated alternative splicing of ETS1, resulting in splice variant P51-ETS1 regulating the expression of tight junction related proteins ZO-1, occludin and claudin-5, that is, regulate the permeability of BTB through the paracellular pathway. The TEER value and HRP flux in this study are important indicators for evaluating the integrity and permeability of BTB. A decrease in TEER value and an increase in HRP flux indicate an increase in BTB permeability^[23]^.

Currently found m^6^A methyltransferases include METTL3, METTL14, WTAP, METTL4 and so on^[24–26]^. This study found that METTL3 is highly expressed in GECs, and the silence of METTL3 increases the permeability of BTB. The results show that METTL3 is involved in the regulation of BTB permeability. In recent years, research on the role of m^6^A modification in various stages of the RNA life cycle has made significant progress^[27]^. M^6^A regulates target mRNAs or miRNAs and participates in the progression of various cancers^[28]^. In a variety of tumor tissues, the upregulation of METTL3 expression plays a carcinogenic role, accompanied by increased m^6^A levels. For example, high levels of METTL3 have been found in various tumors such as liver cancer and lung cancer^[29]^. Through UALCAN (http://ualcan.path.uab.edu) analysis TCGA database data discovery that METTL3 expression increases and the poor prognosis of malignant glioma were positively correlated^[29]^. After silenced METTL3, the detection revealed that BTB permeability was significantly increased, suggesting that METTL3 is involved in the regulation of BTB permeability. To further clarify the role of METTL3, this study revealed that CPEB2 may be a downstream target of METTL3 by MeRIP-Seq data. In order to verify that CPEB2 is the target of METTL3, this study performed m^6^A dot blot assay and MeRIP-qPCR after silencing METTL3, results showed that both overall methylation level and methylation level of CPEB2 decreased significantly, proving that METTL3 promoted the occurrence of CPEB2 m^6^A methylation modification.

The IGF2BP family consists of three members, named IGF2BP1, IGF2BP2 and IGF2BP3^[31]^. Some studies have found that IGF2BP3 is highly expressed in a variety of tumors and is a potential therapeutic target^[32]^, but its function in blood vessels has not been reported yet. This study found that in IGF2BPs, compared with IGF2BP1 and IGF2BP2, the high expression of IGF2BP3 in GECs is statistically significant. After silencing IGF2BP3, the permeability of BTB increased, proving that IGF2BP3 plays a regulatory role in the BTB permeability. In eukaryotes, IGF2BPs are recognized m^6^A readers^[33]^. Related studies have shown that in hepatocellular carcinoma cells, IGF2BPs regulate MYC expression in an m^6^A-dependent manner^[34]^; IGF2BP1 reduces miRNA-mediated down-regulation of SRF mRNA and promotes SRF expression in an m^6^A-dependent manner^[35]^; IGF2BP2 targets DANCR containing m^6^A to enhance its translation, and then IGF2BP2 and DANCR jointly promote the occurrence of pancreatic cancer^[36]^. In this study, RIP and RNA pulldown experiments proved that IGF2BP3 directly recognizes and binds to the m^6^A site on the 3’UTR of CPEB2 mRNA. Through half-life and nascent RNA assays, it is found that IGF2BP3 enhances the stability of CPEB2 mRNA.

This study found that CPEB2 is highly expressed in GECs, and BTB permeability increases after silencing CPEB2. The study results suggest that CPEB2 is involved in the regulation of BTB permeability. Through half-life and nascent RNA assays, it was found that METTL3 enhanced CPEB2 mRNA stability. Similar studies have shown that in gastric cancer cells, METTL3 promotes the m^6^A modification of recombinant protein HDGF mRNA, while IGF2BP3 directly recognizes and binds to the m^6^A site on HDGF mRNA, enhancing the stability of HDGF mRNA^[37]^. Therefore, this study proves that METTL3 promotes CPEB2 mRNA m^6^A modification of CECs, and IGF2BP3 directly recognizes and binds to the m^6^A site on CPEB2 mRNA, enhances the stability of CPEB2 mRNA, and then regulates BTB permeability.

Next, we performed PCR array and qRT-PCR verification on the silenced CPEB2 GECs, and found that the expression level of SRSF5 after CPEB2 silencing decreased significantly, suggesting that SRSF5 may be the target of CPEB2. This study found that SRSF5 is highly expressed in GECs, and BTB permeability increases after silencing SRSF5, suggesting that SRSF5 is involved in the regulation of BTB permeability. This study confirmed the binding effect of CPEB2 and SRSF5 through RIP and RNA pulldown assays, after silencing CPEB2, the expression level of SRSF5 was significantly reduced, the half-life was shortened, but there was no significant change in the level of nascent RNA, suggesting that CPEB2 may regulate BTB permeability by combining SRSF5 mRNA and increasing its stability. SRSF5 is a member of the SR protein family of splicing factors. Studies have shown that in prostate cancer, SRSF1 and SRSF5 are involved in the selective splicing of HSD17B2 mRNA; in lung cancer, SRSF5 regulates tumor growth through AS of CCAR1 pre-mRNA^[37]^. In order to determine that CPEB2 mediates SRSF5 to regulate BTB permeability through m^6^A modification, this study further mutated the CPEB2 m^6^A methylation site and found that the expression level of SRSF5 was significantly reduced and BTB permeability was significantly increased; however, overexpressing CPEB2 while mutating the m^6^A site, it was found that neither the expression level of SRSF5 nor the permeability of BTB had significant changes. The above results indicate that CPEB2 m^6^A methylation promotes CPEB2’s regulation of SRSF5 stability and thus regulates BTB permeability.

Studies have shown that the expression of the transcription factor ETS family is essential for endothelial cell differentiation^[41]^. ETS1 and ETS2 may affect tumor angiogenesis and metastasis in the tumor microenvironment, especially in endothelial cells ^[42]^. In this study, two siRNAs (S1, S2) were designed in exon VII of ETS1, and siRNAs (S3, S4) were designed in exon VI and exon VIII, respectively. Compared with the corresponding si-NC group, si- S1 and si-S2 significantly reduced the mRNA and protein expression of P51-ETS1, while the mRNA and protein expression levels of P42-ETS1 increased significantly; si-S3 and si-S4 significantly reduced the mRNA and protein expression of P51-ETS1 and P42-ETS1, demonstrating that compared to P51-ETS1 mRNA, P42-ETS1 lacks exon VII. It has been reported that P51-ETS1 and P42-ETS1 are different transcription factors in splicing variants of mouse ETS1^[43]^. In MDA-MB-231 breast cancer cells, P51-ETS1 is expressed but there is no P42-ETS1 protein in primary breast cancer, only 10% expressed P42-ETS1 when P51-ETS1 was expressed^[43]^. This study found that in GECs, P51-ETS1 is highly expressed and P42-ETS1 is lowly expressed; silencing P51-ETS1 increases BTB permeability, and overexpressing P51-ETS1 decreases BTB permeability; overexpressing P51-ETS1 can reverses the effect of silencing P51-ETS1 on BTB permeability, while overexpression of P42-ETS1 after silencing P51-ETS1 has no effect on the effect of silencing P51-ETS1, suggesting that P51-ETS1 participates in BTB permeability regulation. The dual-luciferase reporter assays and ChIP assays proved the binding effect of P51-ETS1 with the target genes ZO-1, occludin and claudin-5 promoters. The above results confirm that P51-ETS1 can directly bind to the promoter regions of ZO-1, occludin and claudin-5, promoting their transcription, and then regulating BTB permeability.

In order to study the mechanism of SRSF5 regulating ETS1 alternative splicing, we further silenced SRSF5 and found that the expression of P51-ETS1 was significantly reduced, while the expression level of P42-ETS1 was significantly increased. The analysis of RIP assays and small gene report assays confirmed that in GECs SRSF5 plays a role in inducing ETS1 exon inclusion, causing high expression of P51-ETS1 (exon VII contained) and low expression of P42-ETS1 (exon VII lose); silencing SRSF5 while overexpressing P51-ETS1, reversed the effect of silencing SRSF5 alone on BTB permeability; silencing SRSF5 while overexpressing P42-ETS1 had no significant effect of silencing SRSF5 alone. The above results indicate that SRSF5 enhances the expression level of spliceosome P51-ETS1 by inducing inclusion of ETS1 exon VII, thereby regulating the permeability of BTB. We further silenced CPEB2 and SRSF5, and found that the expression level of P51-ETS1 was significantly reduced, the expression level of P42-ETS1 was significantly increased, and the permeability of BTB was further increased. Overexpression of SRSF5 while silencing CPEB2 reversed the downstream effects of silencing CPEB2 alone. The above results indicate that CPEB2 and SRSF5 are jointly involved in regulating BTB permeability.The major alternative splicing patterns include exon skipping, intron retention, mutually exclusive exons and alternative 3′ or 5′ splice sites, which forms through the recognition of splicing regulatory elements (SREs)^[45]^. SREs recruit proteins and complexes that enhance or silence splicing and have been named descriptively: ESEs, intronic splicing enhancers (ISEs), exonic and intronic splicing silencers (ESSs and ISSs)^[46]^. ESEs can increase the exon inclusion body by acting as the binding site for the assembly of multicomponent splicing enhancer complex, which is generally recognized by at least one member of SR family^[47]^. SR protein is widely regarded as a positive splicing regulator and can promote exon inclusion by binding with exonic splicing enhancers (ESEs)^[48]^. SRSF1 can promote the exon inclusion of CD33 in Alzheimer’s disease, enhance the transcriptional expression of full-length CD33, and regulate their specific interaction with CD33 pre-mRNA to change the protein level on the cell surface^[49]^; SRSF6 increased the inclusion of OGDHL exon 3, further affecting pancreatic cancer cell metastasis^[50]^. The mechanism by which SRSF5 promotes the inclusion of ETS1 exon VII in GECs and thereby regulates the permeability of BTB was found in this study to enrich the function of SR protein in alternative splicing.

Dox is an anthracycline antitumor antibiotic, which is used in the clinical treatment of cancer^[51]^, but for the treatment of brain tumors, due to the presence of the blood-brain barrier, it is difficult for Dox to enter the brain parenchyma and reach an effective therapeutic concentration^[52]^. Studies have confirmed that the combined application of KHDRBS3 and doxorubicin can promote the transmembrane transport of doxorubicin and induce apoptosis of glioma cells^[53]^. In order to further evaluate the regulatory effect of the above regulatory factors on BTB permeability in this study, we combined Dox with the BTB model established by the above regulatory factors to observe the effect on the apoptosis of U251 glioma cells. The results showed that the silencing of METTL3, IGF2BP3, CPEB2, SRSF5 and ETS1 alone, as well as CPEB2, SRSF5, P51-ETS1 triple silenced, all significantly increased the rate of Dox-induced apoptosis of U251 glioma cells. The results suggest that the application of METTL3, IGF2BP3, CPEB2, SRSF5 and P51-ETS1 alone can enhance the effect of Dox-induced apoptosis of glioma cells. When CPEB2, SRSF5 and P51-ETS1 were used in combination, the effect of inducing apoptosis of glioma cells was statistically significant compared with that of single application.

## Conclusions

In summary, the upregulated METTL3 and IGF2BP3 in GECs increase the mRNA stability of CPEB2 mRNA by m^6^A methylation and upregulate the expression of CPEB2; CPEB2 binds and increases the stability of the splicing factor SRSF5 mRNA; SRSF5 promotes the appearance of ETS1, the spliceosome P51-ETS1 promotes the transcriptional expression of tight junction related proteins ZO-1, occludin and claudin-5, thereby regulating the BTB permeability. The use of METTL3, IGF2BP3, CPEB2, SRSF5, and P51-ETS1 alone or in combination can effectively enhance Dox through BTB and thus promote the apoptosis of glioma cells. The results of this study not only provide a new theoretical and experimental basis for the molecular regulation of BTB from the perspective of epigenetics, but also provide a new idea for the comprehensive treatment of glioma.

## Legend

### 1. METTL3 is highly expressed in GECs, knockdown METTL3 increases BTB permeability

**Fig. S1.**
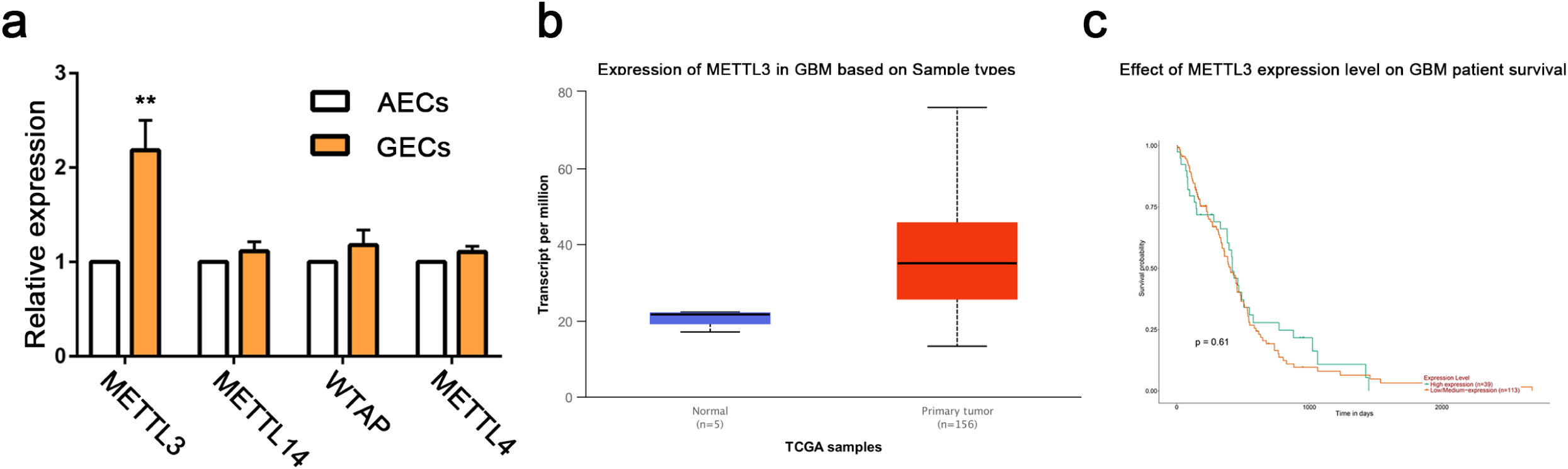
Measurement of METTL3 expression level in GECs and analysis of expression level in Glioblastoma multiforme and its influence on prognosis. **a** Relative mRNA levels of METTL3, METTL14,WTAP and METTL4 in AECs and GECs were determined by qRT-PCR. Data represented as mean ± SD (n = 3). \*\**P* < 0.01 vs. AECs group. **b** Expression of METTL3 in Glioblastoma multiforme (GBM) based on sample types by UALCAN. **c** Effect of METTL3 expression level on GBM patient survival by UALCAN.

### 2. METTL3 mediated m6A methylation of CPEB2 mRNA

**Fig. S2.**
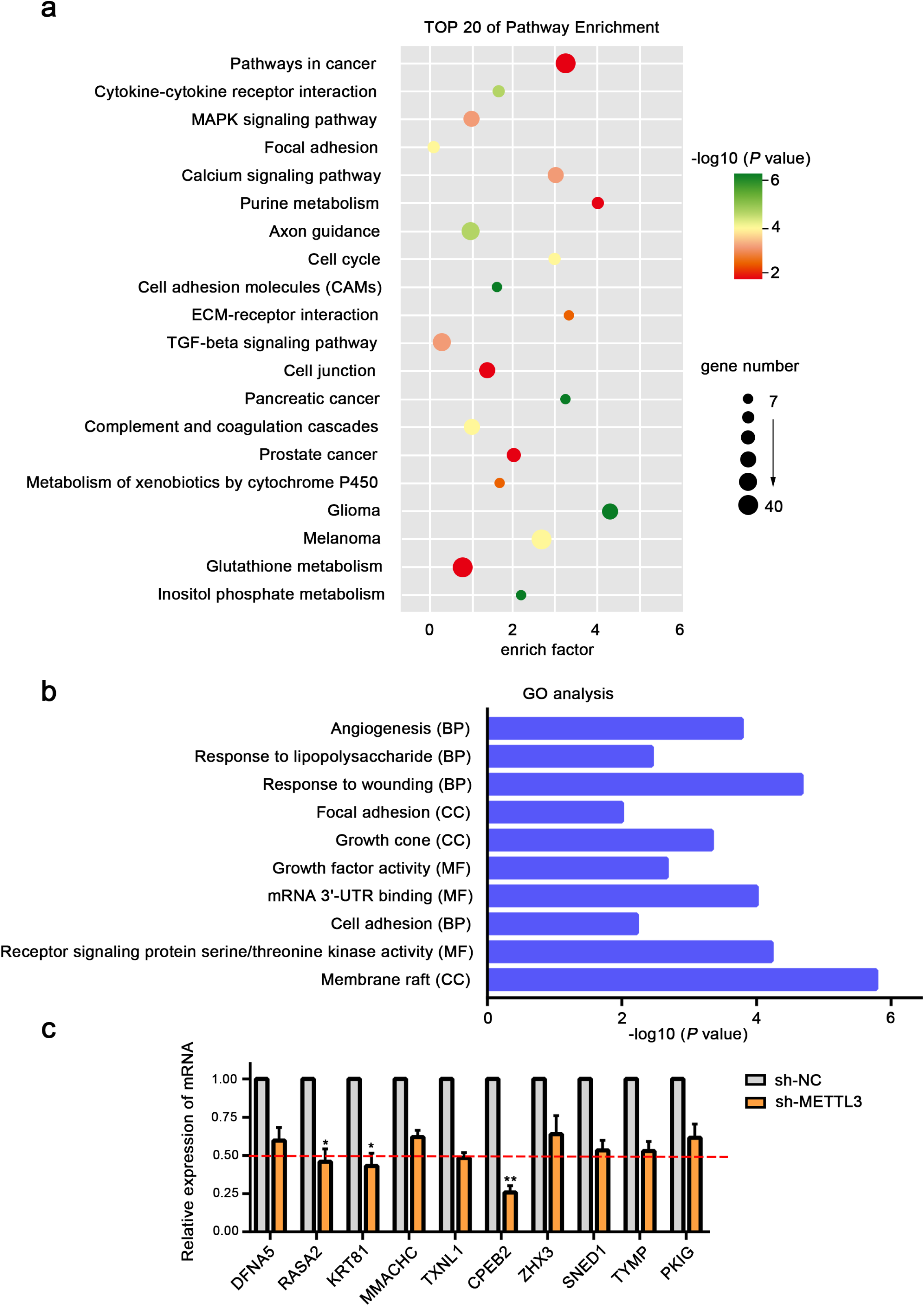
Screening of target genes. **a-b** Pathway enrichment and GO analyze. **c** Relative mRNA levels of ten differential genes in sh-NC and sh-METTL3 in GECs were determined by qRT-PCR. Data represented as mean ± SD (n = 3). *P < 0.05, **P < 0.01 vs. sh-NC group.

### 3. CPEB2 is highly expressed in GECs, silencing CPEB2 increases BTB permeability; IGF2BP3 combined with CPEB2 mRNA 3’-UTR increases its mRNA stability

**Fig S3.**
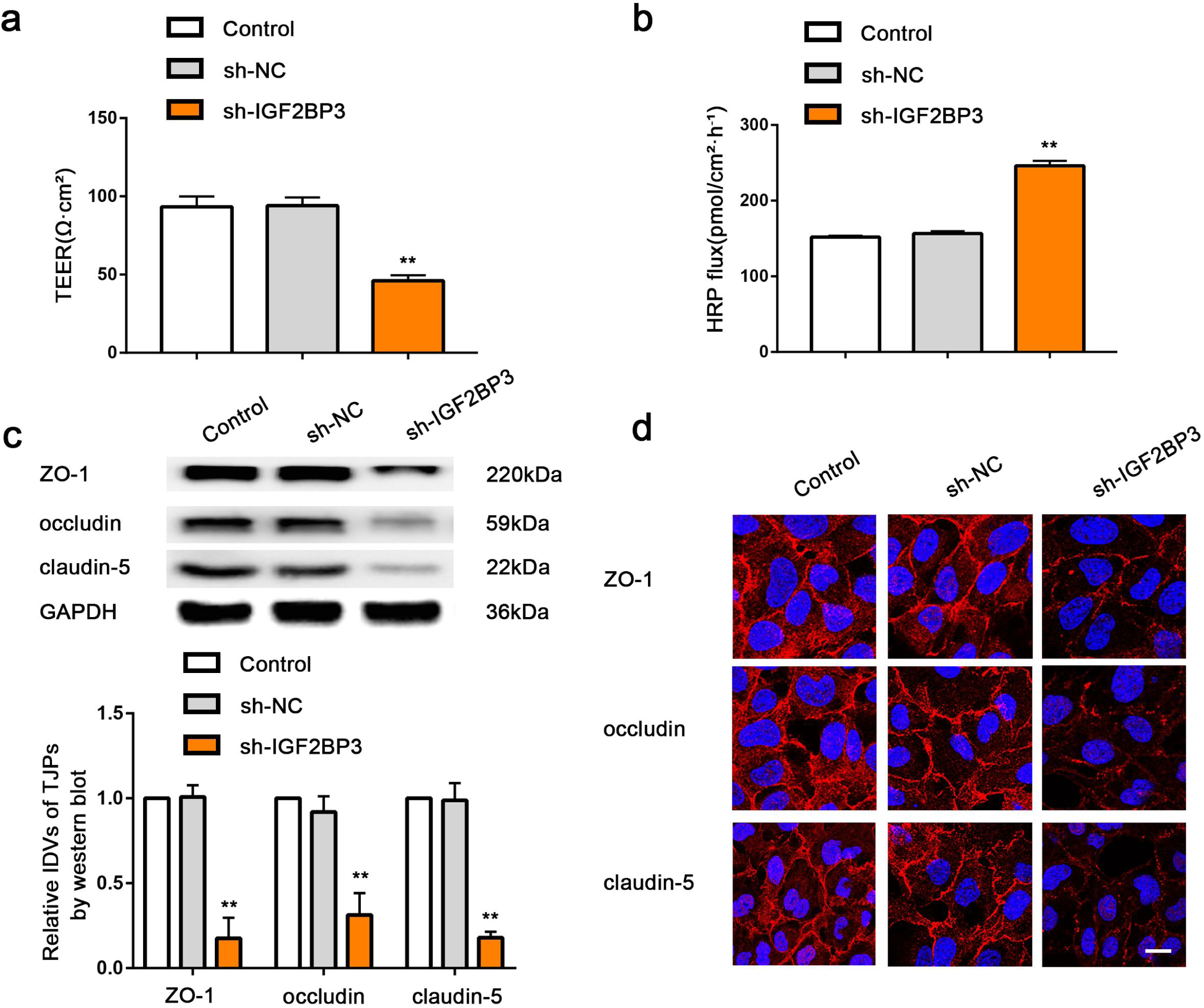
METTL3 interacted with IGF2BP3 to regulate BTB permeability. a-b. Effects of METTL3 and IGF2BP3 double knockdown on TEER values and HRP flux *in vitro* BTB model. **c** Effects of METTL3 and IGF2BP3 double knockdown on the expressions of ZO-1, occludin and claudin-5 were analyzed by western blot. Data represented as mean ± SD (n = 3). \*\**P* < 0.01 vs. sh-METTL3-NC group. **d** The distributions of ZO-1, occludin and claudin-5 in the METTL3 and IGF2BP3 double knockdown GECs were observed by immunofluorescent staining. Scale bar represents 50 µm.

### 4. SRSF5 is highly expressed in GECs, and silencing SRSF5 increases the permeability of BTB; CPEB2 enhances the stability of SRSF5 mRNA and regulates the permeability of BTB

**Fig.S4.**
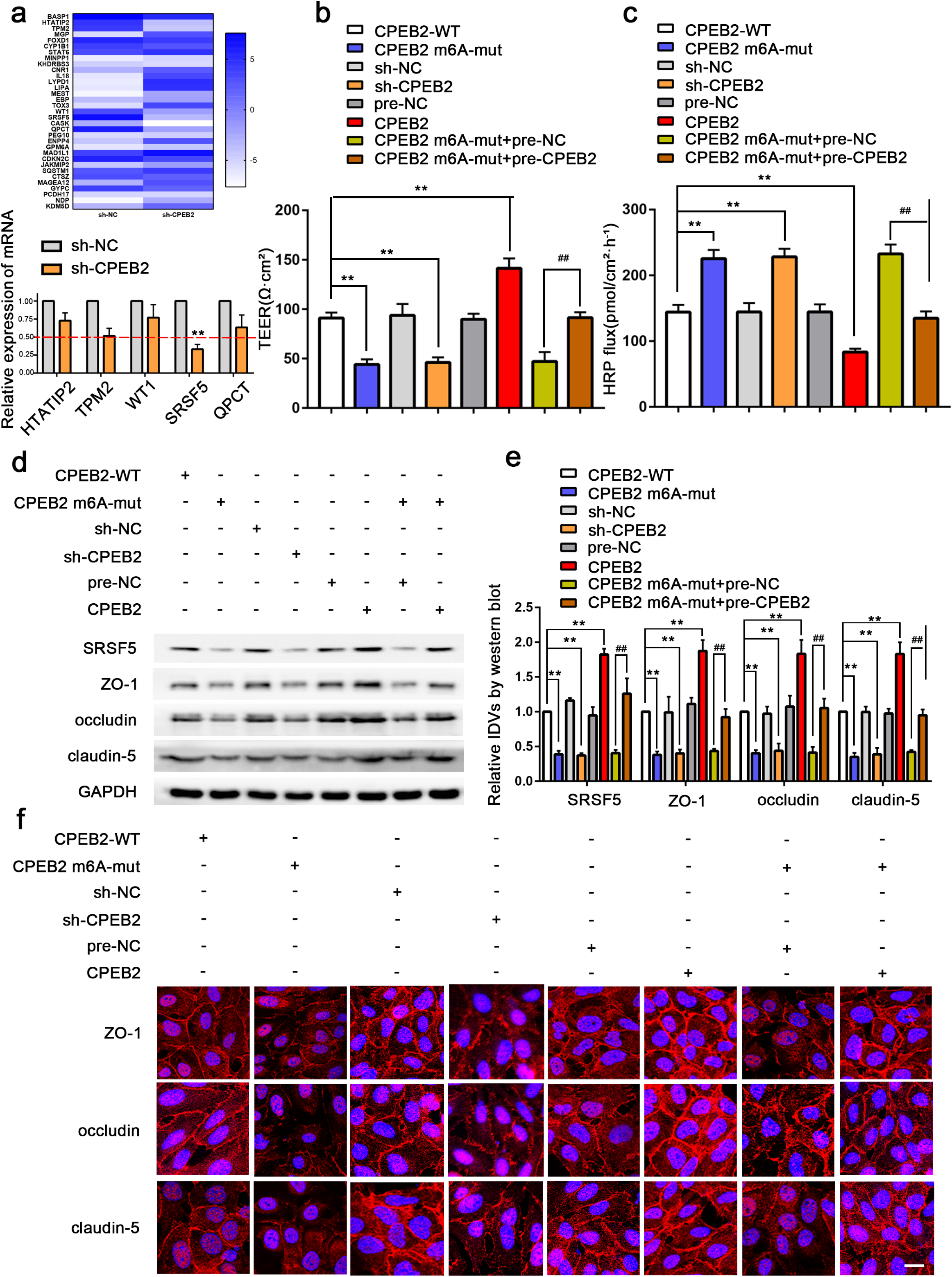
CPEB2 interacted with SRSF5 to regulate BTB permeability. **a** PCR array for sh-CPEB2 and qRT-PCR to detect differential genes, \*\**P* < 0.01 vs. AECs group. **b-c** Effects of CPEB2 m^6^A-mut, sh-CPEB2, pre-CPEB2 and CPEB2 m^6^A-mut+pre-CPEB2 on TEER values and HRP flux *in vitro* BTB model. **d-e** Effects of CPEB2 m^6^A-mut, sh-CPEB2, pre-CPEB2 and CPEB2 m^6^A-mut +pre-CPEB2 on the expressions of ZO-1, occludin and claudin-5 were analyzed by western blot. Data represented as mean ± SD (n = 3). \*\**P* < 0.01 vs. CPEB2-wt group, ##*P* < 0.01 vs. CPEB2 m^6^A-mut+pre-NC group. **f** The distributions of ZO-1, occludin and claudin-5 in the CPEB2 m^6^A-mut, sh-CPEB2, pre-CPEB2 and CPEB2 m^6^A-mut+pre-CPEB2 GECs were observed by immunofluorescent staining. Scale bar represents 50 µm.

### 5. P51-ETS1 is highly expressed in GECs, silencing P51-ETS1 increases the permeability of BTB

**Fig.S5 Analysis.**
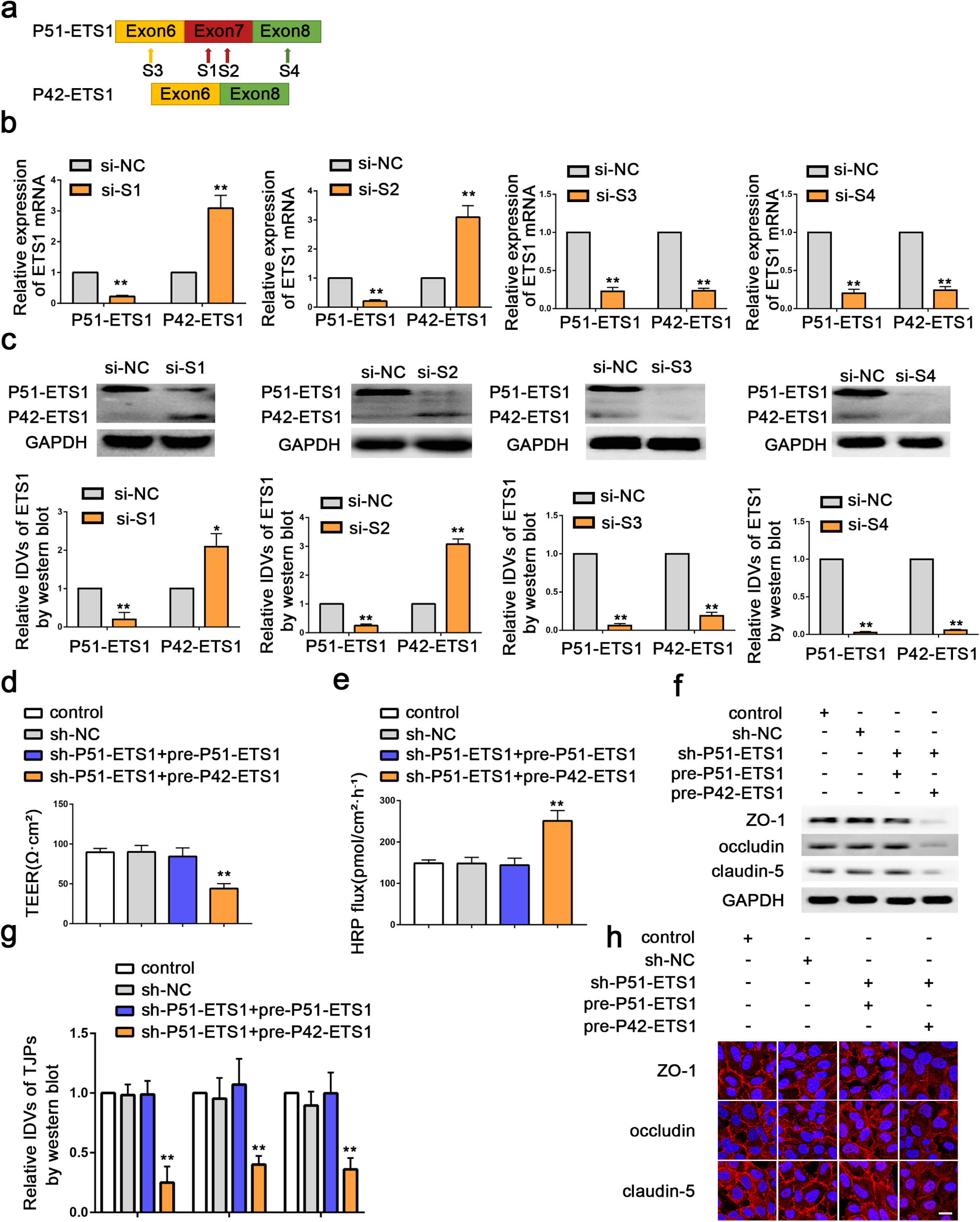
of ETS1 splicing variants. **a** The model of two pairs of specific small interfering RNAs (siRNAs),one pair (si-S1 and si-S2) for knocking down ETS1 exon7 inclusion P51-ETS1 variants, another pair (si-S3 and si-S4) for knocking down both ETS1 exon7 inclusion P51-ETS1 and ETS1 exon7 skipping P42-ETS1 variants. **b,c** qRT-PCR and western blot for detecting P51-ETS1 and P42-ETS1 splicing isoforms expression in GECs with si-S1, si-S2, si-S3 and si-S4. Data represented as mean ± SD (n = 3). \**P* < 0.05, \*\**P* < 0.01 vs. si-NC group. **d-e** Effects of sh-P51-ETS1, sh-P51-ETS1+pre-P51-ETS1, sh-P51-ETS1+pre-P42-ETS1 on TEER values and HRP flux *in vitro* BTB model. **f-g** Effects of sh-P51-ETS1, sh-P51-ETS1+pre-P51-ETS1,sh-P51-ETS1+pre-P42-ETS1 on the expressions of ZO-1, occludin and claudin-5 were analyzed by western blot. Data represented as mean ± SD (n = 3). \*\**P* < 0.01 vs. sh-NC group. **h** The distributions of ZO-1, occludin and claudin-5 in the sh-P51-ETS1, sh-P51-ETS1+pre-P51-ETS1, sh-P51-ETS1+pre-P42-ETS1 GECs were observed by immunofluorescent staining. Scale bar represents 50 µm.

### 6. SRSF5 induces ETS1 exon VII inclusion, and P51-ETS1 binds to the TJPs promoter region

**Fig.S6.**
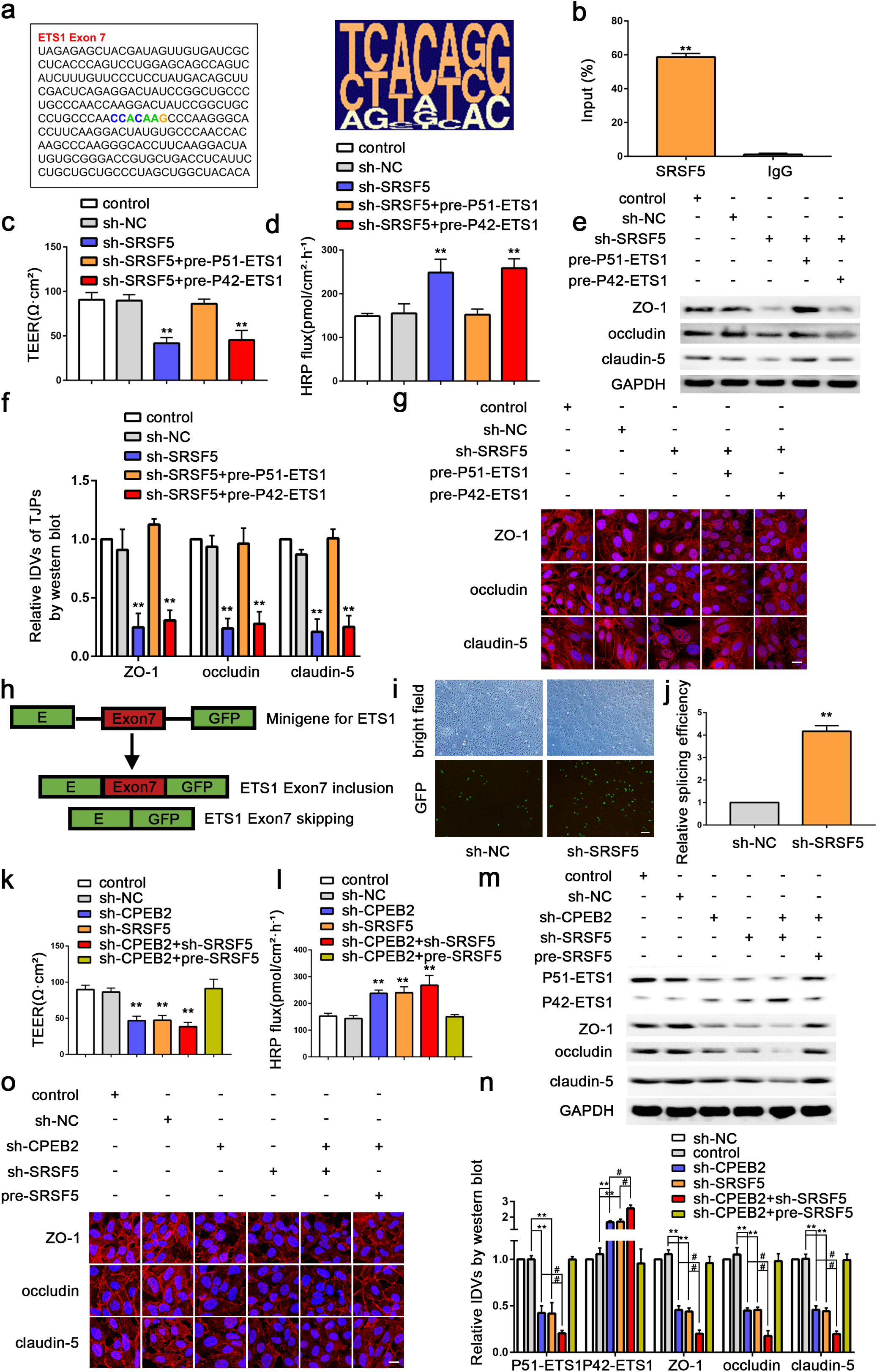
SRSF5 induces ETS1 exon inclusion. **a** A consensus motif sequence (CCACAAG) with predicted SRSF5 binding motif in exon7 of ETS1. **b** RIP for detecting the interaction between SRSF5 protein and ETS1 mRNA. **c,d** Effects of sh-SRSF5, sh-SRSF5+pre-P51-ETS1, sh-SRSF5+pre-P42-ETS1 on TEER values and HRP flux *in vitro* BTB model. **e**,**f** Effects of sh-SRSF5, sh-SRSF5+pre-P51-ETS1, sh-SRSF5+pre-P42-ETS1 on the expressions of ZO-1, occludin and claudin-5 were analyzed by western blot. Data represented as mean ± SD (n = 3). \*\**P* < 0.01 vs. sh-NC group. **g** The distributions of ZO-1, occludin and claudin-5 in the sh-SRSF5, sh-SRSF5+pre-P51-ETS1, sh-SRSF5+pre-P42-ETS1 GECs were observed by immunofluorescent staining.Scale bar represents 50 µm. **h** Minigene reporter system for detecting ETS1 exon7 splicing. **i** The fluorescence signal in the GFP channel represents exon7 splicing efficiency in sh-NC and sh-SRSF5 GECs.Scale bar represents 200 µm. **j** Quantification of splicing efficiency by measuring the relative expression of intact EGFP transcript ETS1 mRNA levels. **k,l** Effects of sh-CPEB2, sh-SRSF5, sh-CPEB2+sh-SRSF5, sh-CPEB2+pre-SRSF5 on TEER values and HRP flux *in vitro* BTB model. **m,n** Effects of sh-CPEB2, sh-SRSF5, sh-CPEB2+sh-SRSF5, sh-CPEB2+pre-SRSF5 on the expressions of ZO-1, occludin and claudin-5 were analyzed by western blot. Data represented as mean ± SD (n = 3). \*\**P* < 0.01 vs. sh-NC group. **o** The distributions of ZO-1, occludin and claudin-5 in the sh-CPEB2, sh-SRSF5, sh-CPEB2+sh-SRSF5, sh-CPEB2+pre-SRSF5 GECs were observed by immunofluorescent staining. Scale bar represents 50 µm.

### 7. The combined application of multiple targets promotes doxorubicin (Dox) to induce glioma cell apoptosis

## Acknowledgments

This work is supported by grants from the Natural Science Foundation of China (81872503, 81872073); China Postdoctoral Science Foundation (2019M661172); Liaoning Science and Technology Plan Project (2020-BS-097and 2017225020).

## Authors’ contributions

YX.X. and YH.L. contributed to conceived and designed the project. MY.Z., CQ.Y., XB.L., and XL.R. contributed to performed most of the experiments. H.C., J.Z., LQ.S. and D.W. did the statistics analysis. MY.Z., CQ. Y. and XB.L. wrote the manuscript.

P.W. Z. L. and B.Y. did western blotting experiments. YX.X. and YH.L. contributed to the manuscript revision. All authors read and approved the fifinal manuscript.

---

**Table S1.**
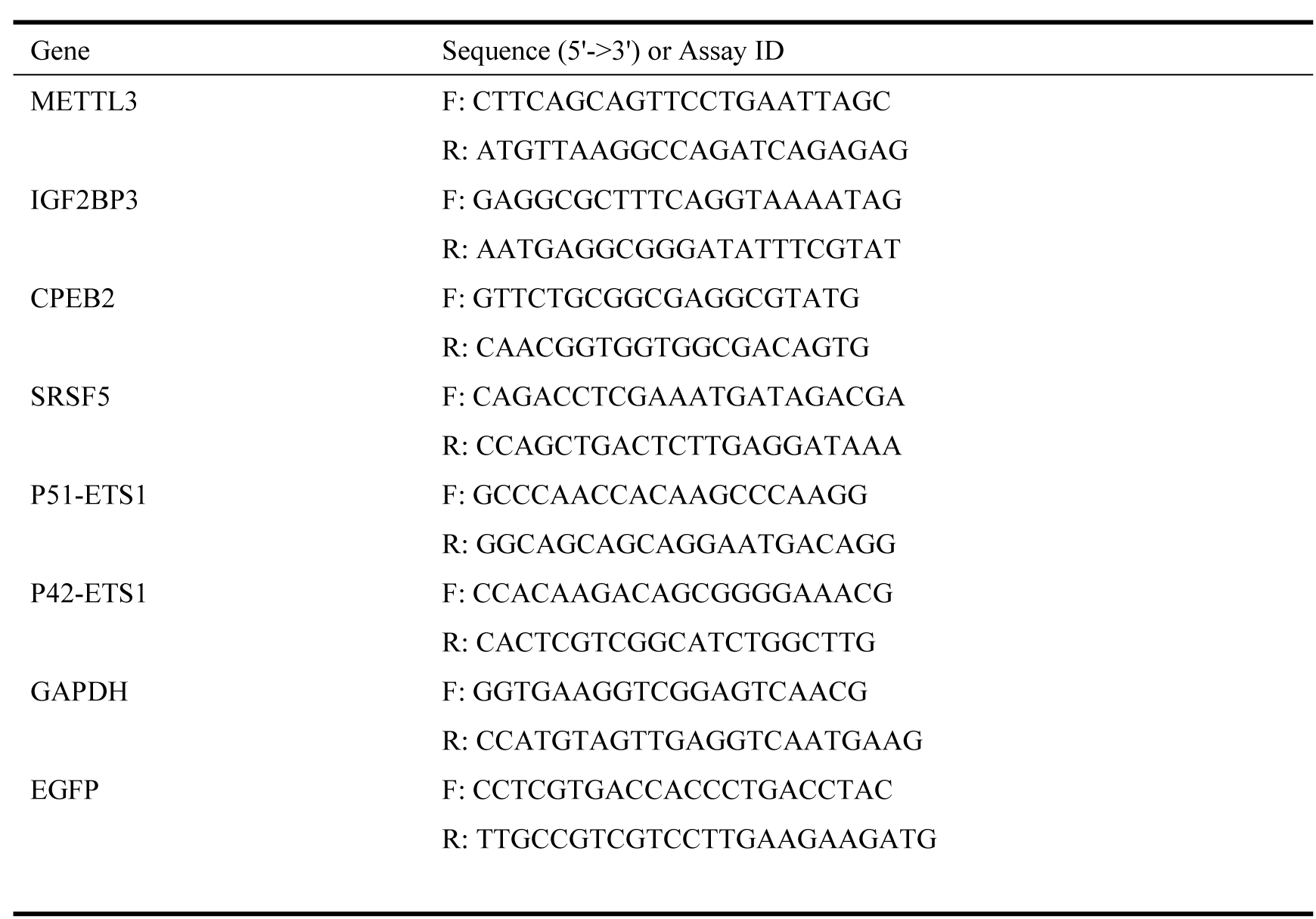
Primers used for qRT-PCR

**Table S2.**
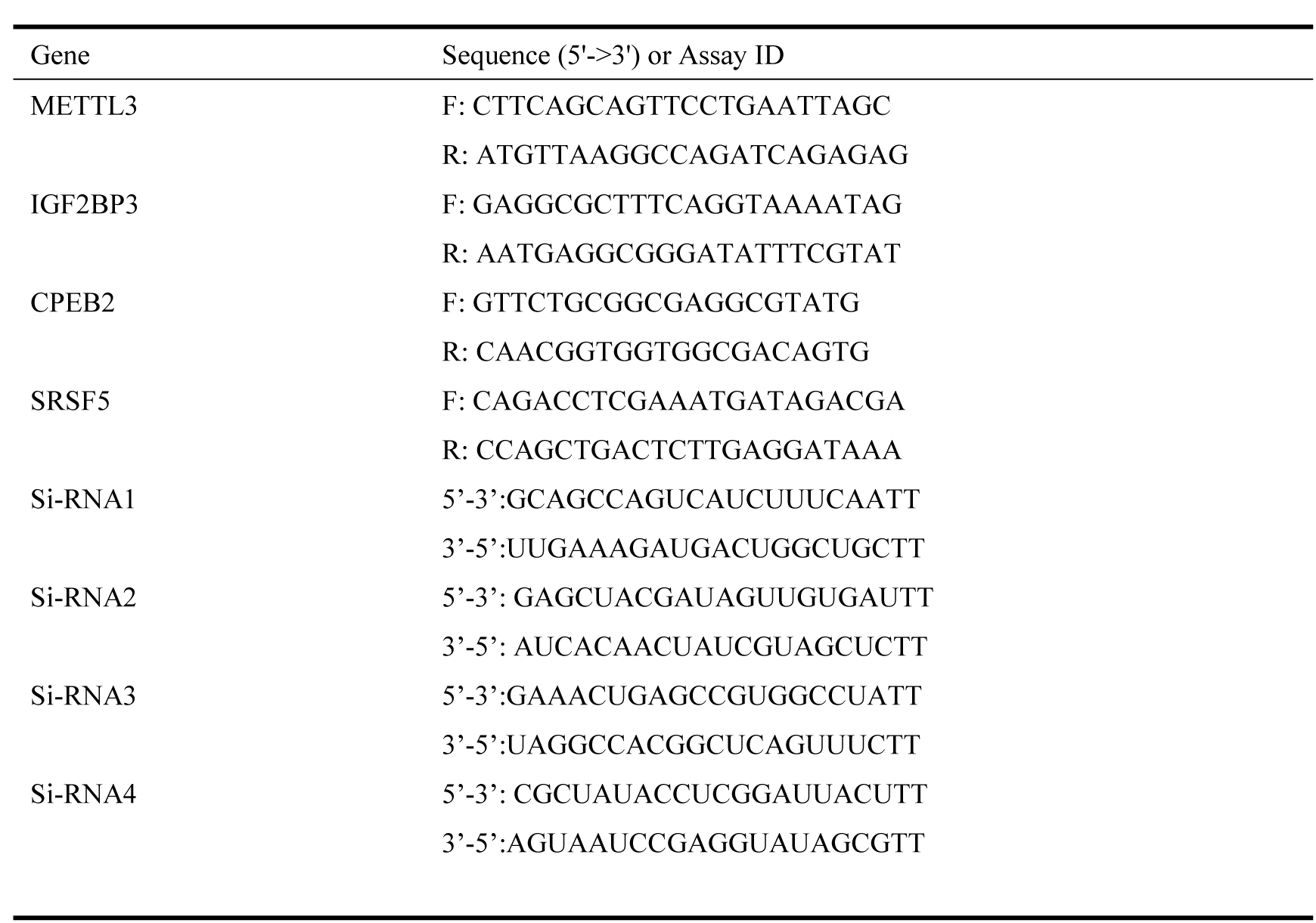
shRNA.

**Table S3.** Antibodies used for western blotting (WB), RNA-binding protein immunoprecipitation (RIP)

| **Protein** | **Applications** | **Antibody** | **Origin** | **Dilution** | **Molecular weight** |
| --- | --- | --- | --- | --- | --- |
| METTL3 | WB | 1507-1-AP, proteintech | Rabbit | 1:500 | 64kD |
| IGF2BP3 | WB,IP | 14642-1-AP, proteintech | Rabbit | 1:500;1µg per IP | 66kD |
| CPEB2 | WB,IP,IF | ab51069, Abcam | Rabbit | 1:500;1µg per IP;1:50 | 66kD |
| SRSF5 | WB,IP | Ab67175, Abcam | Mouse | 1:500;1µg per IP | 65kD |
| P51-ETS1 | WB | 12118-1-AP, proteintech | Rabbit | 1:500 | 51kD |
| P51-ETS1 | IP | Sc-55581, Santa | Mouse | 1µg per IP | 51kD |
| P42-ETS1 | WB | 12118-1-AP, proteintech | Rabbit | 1:500 | 42kD |
| GAPDH | WB | 60004-1-Ig, proteintech | Mouse | 1:10000 | 36kD |
| Ago2 | RIP | 03-110, MerckMillipore | Mouse | 1:10 | 100kD |
| IgG | RIP | ab18413, Abcam | Mouse | 1:10 | 150kD |
| ZO-1 | WB,IF | 61-7300, Thermo Fisher | Rabbit | 1:500;1:50 | 220kD |
| Occludin | WB,IF | 71-1500, Thermo Fisher | Rabbit | 1:500;1:50 | 59kD |
| Claudin-5 | WB,IF | 35-2500, Thermo Fisher | Mouse | 1:500;1:50 | 22kD |
| Goat anti-mouse | WB | SA00001-1, proteintech |  | 1:10000 |  |
| Goat anti-rabbit | WB | SA00001-2, proteintech |  | 1:10000 |  |

**Table S4.**
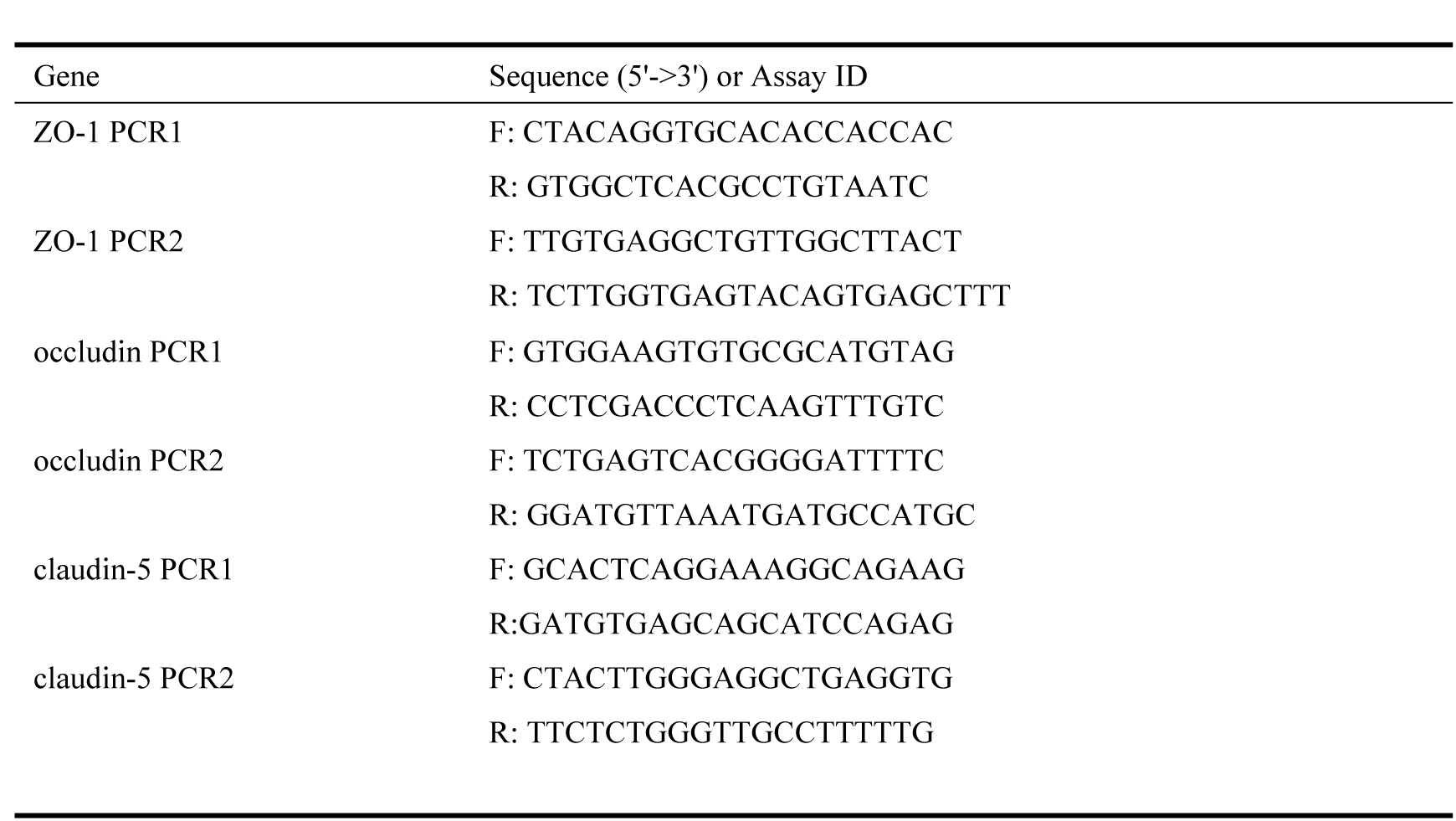
Primers used for ChIP.

**Table S5.**
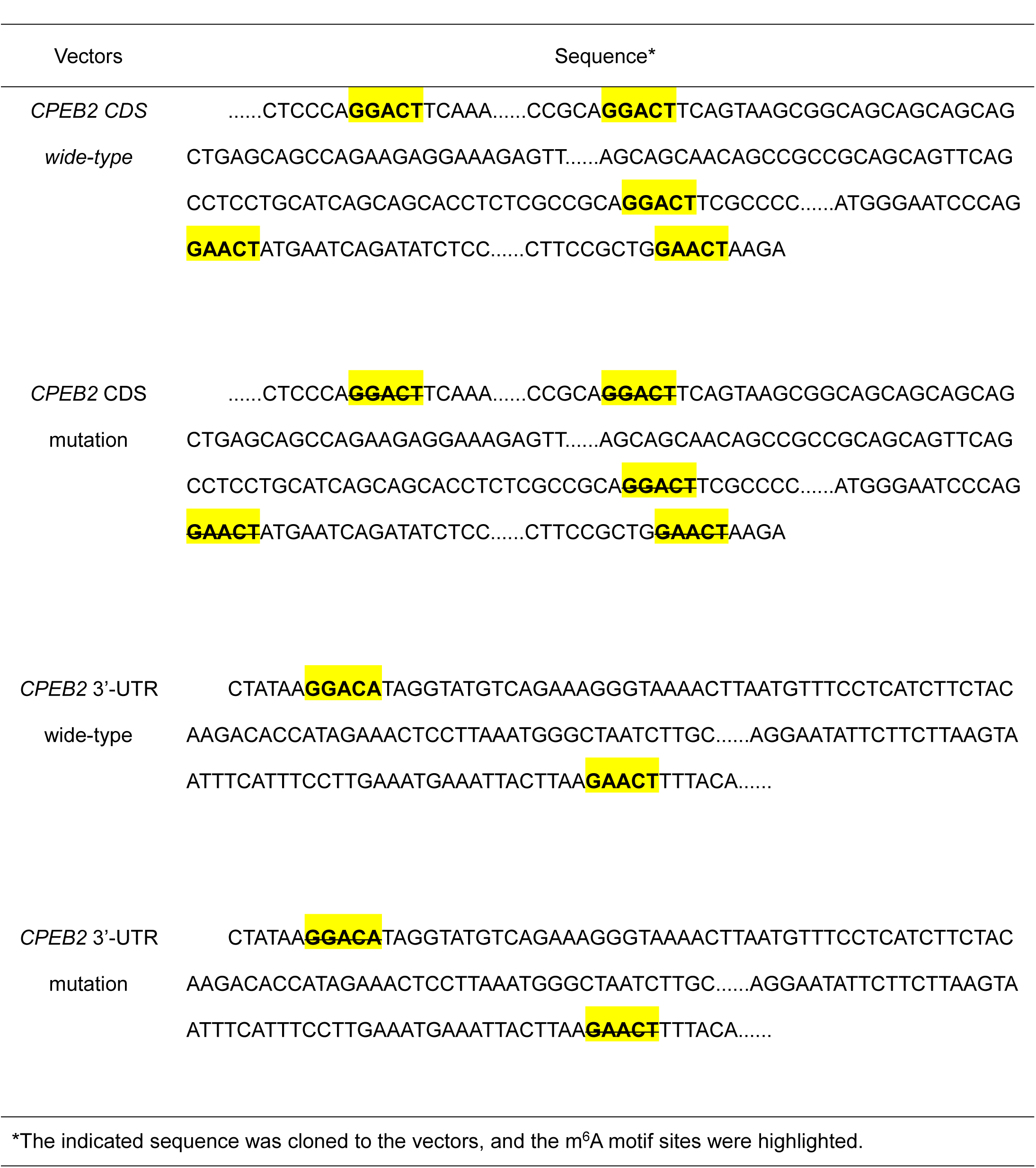
The specific sequence of wide-type or m^6^A motif depletion *CPEB2* CDS and 3’-UTR.

